# Social interaction and network structure in groups of *Drosophila* males are shaped by prior social experience and group composition

**DOI:** 10.1101/2020.03.19.995837

**Authors:** Assa Bentzur, Shir Ben-Shaanan, Jennifer Benishou, Eliezer Costi, Amiyaal Ilany, Galit Shohat-Ophir

## Abstract

Living in a group creates a complex and dynamic environment in which the behavior of the individual is influenced by and affects the behavior of others. Although social interactions and group living are fundamental adaptations exhibited by many organisms, relatively little is known about how prior social experience, internal states and group composition shape behavior in a group, and the neuronal and molecular mechanisms that mediate it. Here we present a practical framework for studying the interplay between social experience and group interaction in *Drosophila melanogaster* and show that the structure of social networks and group interactions are sensitive to group composition and individuals’ social experience. We simplified the complexity of interactions in a group using a series of experiments in which we controlled the social experience and motivational states of individuals to dissect patterns that represent distinct structures and behavioral responses of groups under different social conditions. Using high-resolution data capture, machine learning and graph theory, we analyzed 60 distinct behavioral and social network features, generating a comprehensive representation (“group signature”) for each condition. We show that social enrichment promotes the formation of a distinct group structure that is characterized by high network modularity, high inter-individual and inter-group variance, high inter-individual coordination, and stable social clusters. Using environmental and genetic manipulations, we show that this structure requires visual and pheromonal cues, and that cVA sensing neurons are necessary for the expression of different aspects of social interaction in a group. Finally, we explored the formation of group behavior and structure in heterogenous groups composed of flies with distinct internal states, and discovered evidence suggesting that group structure and dynamics reflect a level of complexity that cannot be explained as a simple average of the individuals that constitute it. Our results demonstrate that fruit flies exhibit complex and dynamic social structures that are modulated by the experience and composition of different individuals within the group. This paves the path for using simple model organisms to dissect the neurobiology of behavior in complex social environments.

## Introduction

Many species have adapted to living in groups, from simple organisms, such as nematodes, to humans. Group living takes different forms with various levels of complexity, from almost random interactions to fully synchronized collective behavior^1–5^, and can be described by measuring the behavior of individuals, the interaction between individuals and the resulting social network, altogether defined here as “group behavior”. When individuals interact in a group, their previous experience, motivation and physiological state (termed here as internal state) affect their action selection, giving rise to diverse activity levels, behavioral responses, and engagement with others^6–8^. This results in a highly complex and ever changing environment, where each interaction can change the social context of subsequent interactions, leading to a variety of behavioral outcomes from what seem to be identical starting conditions^7,9^. The complex nature of this environment imposes conceptual challenges in the quantification and analysis of group behavior^10^.

A fundamental question in this respect is how internal and external factors such as previous social experience, specific group composition or the existence of available resources, shape group behavior^11,12^. Although much is known about the interplay between social experience, internal states^13–18^ and their effects on social interaction in pairs of animals^14,19–23^, relatively little is known about how these elements shape social behavior in a group. Currently, group behavior is mainly studied at two organizational levels: the behavioral repertoires of individuals within groups, and the structure and dynamics of all interactions within a group (social network analysis)^24^. Both lines of study progressed substantially with advances in machine vision and machine learning technologies that allow automated tracking and unbiased behavioral analysis^25–31^. Analyzing the behavioral repertoires of individuals within a group can provide a comprehensive description of behavioral responses of all individuals under different conditions, enabling the dissection of mechanisms that shape each behavior, the sensory requirements for a given behavior and the specific context it is presented in. However, this approach does not provide much information about group structure. By evaluating every interaction between pairs of individuals in a group, network analysis can be used to represent integrated systems such as social groups, providing insights into the formation, dynamics, and function of group structure^24,32–34^. This type of analysis can be employed to investigate transmission processes in groups as a basis for understanding complex phenomena such as microbe transmission, social grooming, decision making, and hierarchy^3,32,35–47^. Although analysis of individual behaviors and social networks highlight different aspects of social interaction, they are complementary for understanding complex emergent phenomena such as group behavior.

Studies of social interaction in *Drosophila melanogaster* have mainly focused on understanding the neuronal basis of innate and recognizable behaviors such as male–male aggression and male–female courtship encounters^48–53^. Various studies provided mechanistic understanding of these complex behaviors, demonstrating that their expression requires multi-sensory inputs, as well as specific neuronal pathways in the brain^52,54–59^. Modulation of behavior by previous social experience was also investigated in flies, revealing that gene regulation in specific neuronal populations can lead to long-lasting behavioral changes^20,60–64^. The social behavior of *D. melanogaster* in the wild remains largely understudied. Nonetheless, it was shown that wild flies are relatively stationary, moving only a few meters a day, tending to group with conspecifics while avoiding flies of different species^65^. These aggregations seem to be plastic and dynamic and facilitate mating with members of other groups to decrease inbreeding. Aggregations are a substrate for a rich repertoire of social interaction that includes courtship, competition over mating partners, mating and communal oviposition^65^. Sex-specific adaptations for space-use were suggested, possibly driven by avoidance of predators, parasites, or males^66^.

While *Drosophila* proves to be a useful model organism for mechanistic dissection of complex behaviors^67,68^, only a small number of studies examined social interaction in groups of flies. These studies demonstrated that flies possess the neuronal ability to recognize different individuals in a group^69^, that groups of flies exhibit non-random group structures which depend on certain sensory systems^4,59,70^ and group size^71^, and that group interaction facilitates collective responses to threats^4,72^. These findings, together with the existence of dedicated circuits for processing social information, and evidence for the presence of social aggregates in wild flies, support the notion that group living is a fundamental component of *Drosophila* behavior. Still, little is known about how group behavior in *Drosophila* unfolds under different biological and environmental conditions. Specifically, it is not clear whether flies form groups with different structures under various conditions, whether the group is affected by internal properties of its constituting individuals and their composition, by different environmental conditions, and whether individual recognition plays a role in such groups.

To bridge these gaps, we searched for conditions that can facilitate the formation of distinct group behaviors. We hypothesized that groups composed of flies with different social histories such as flies that were socially raised and flies that were socially isolated, will exhibit distinct emergent group structures that result from differences in motivation, experience, activity level, and/or sensory sensitivity of the interacting flies. To analyze the emergent group properties, we established an experimental framework that clusters various behavioral and social network parameters into behavioral “group signatures”. We presumed the group signature of socially raised flies to reflect a snapshot of established relationships between members of the group that developed over the course of the experience phase, while that of solitary flies to reflect initial interaction of flies that are exposed for the first time to other flies. Additionally, studies from various animal species^21,22,73–75^ including *Drosophila* have shown that isolation results in increased activity/arousal, increased aggression and in some cases social avoidance. Extending these findings to group context, we predicted groups of solitary flies to exhibit increased activity, increased aggression and reduced social interaction. In contrast, groups of socially raised flies were predicted to show increased social interaction due to reduced aggression^76,77^. Here we show that social experience can drive the formation of groups with distinct behavior and network structures, and that group signature is a useful tool for simplifying the analysis of the multifaceted repertoire of parameters associated with social interaction in groups. Moreover, we show that the group signature of socially raised flies is strongly influenced by both visual cues and the sensing of the male-specific pheromone 11-cis-vaccenyl acetate (cVA). Finally, we explored social interactions in heterogenous groups and identified clusters of features that are sensitive to increasing ratios of aggressive flies, some of which reveal that inter-individual coordination depends on group composition.

## Results

### Establishing a data capture and analysis pipeline for studying complex behavior in groups

To explore the interplay between social history, internal states and social group interaction, we exposed male flies to distinct social conditions and recorded their social interactions within circular arenas that are suitable for analyzing complex group behavior (Fly Bowl system)^78^. To quantify and analyze the behavioral repertoire of individual flies, group interaction, and the resulting social networks, we adapted the Fly Bowl suite of tracking and behavior analysis tools (Ctrax, JAABA, and JAABA plot, Fig. 1A)^78–80^. Although Ctrax is successfully used in many behavioral setups its output includes some tracking errors such as unifying identities and failure to recognize a fly for several frames, impeding analysis that requires accurate and stable identities throughout the experiment. To resolve this, we developed a secondary processing algorithm for Ctrax output data, named FixTRAX. FixTRAX uses a set of rules to find tracking errors, calculates statistical scores that determine which identities to correct per frame, and generates a graphical summary of tracking quality per movie (detailed explanation of the algorithm, error rate and code are found in the methods section and supplementary FixTRAX files). Corrected output data are used to calculate kinetic features and classify eight distinct complex behaviors using the supervised machine learning algorithm JAABA^78^ (Fig. 1A; full description in Supplementary Table S1).

**Figure 1.**
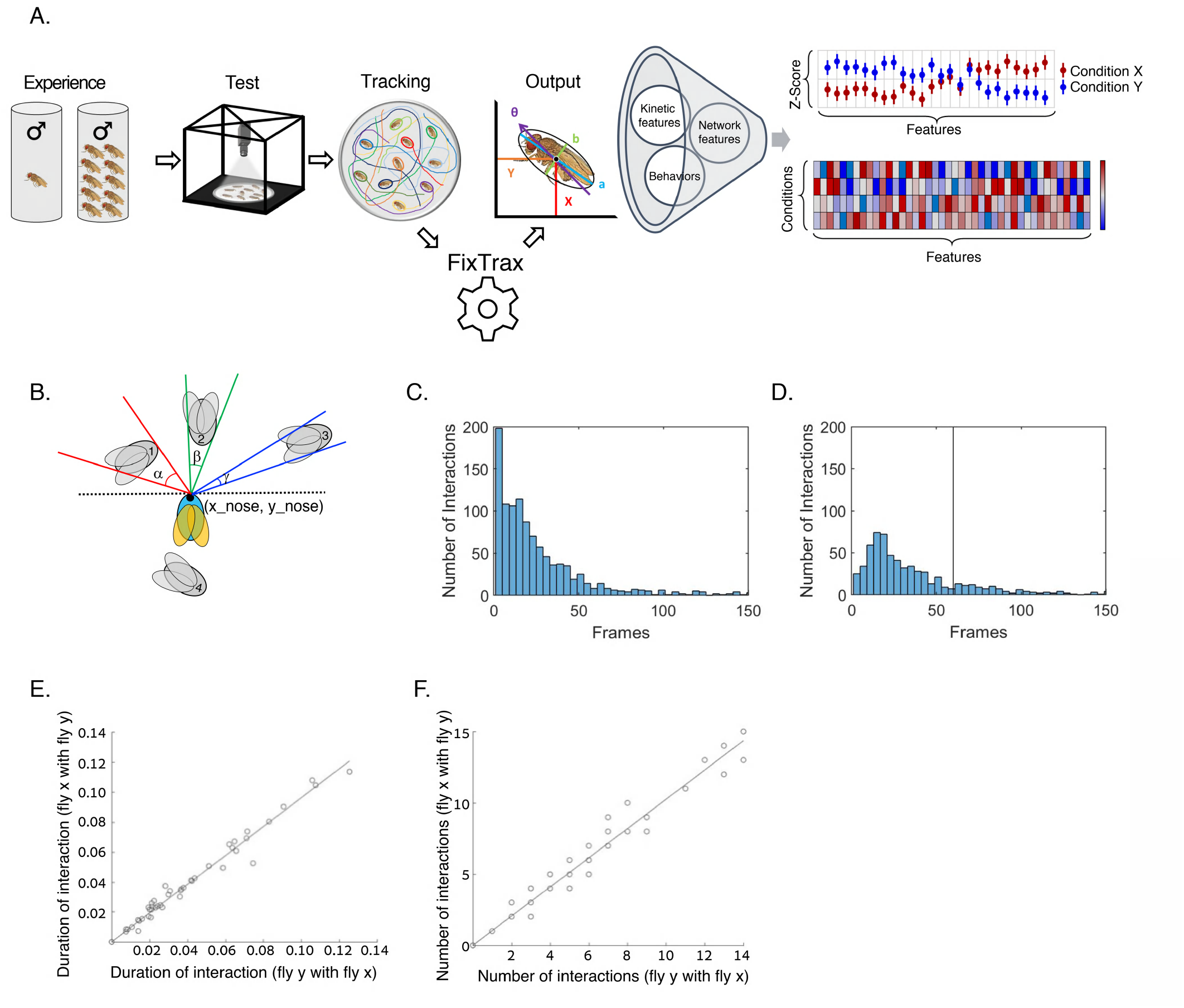
A conceptual and experimental setup for studying complex behavior in groups of *Drosophila*. A. Illustration of social conditioning, data capture and analysis. Naïve male flies were housed in groups of 10 flies or in isolation for 3 days and inserted in groups of 10 into Fly Bowl arenas, where their social interaction was recorded for 15 minutes (at 30fps). Tracking was performed using Ctrax. Error correction of Ctrax output data was performed using FixTRAX, generating an output file of position, angle and size per-fly per-frame. The fixed output file was used to calculate kinetic features, to classify specific behaviors using JAABA and to analyze social network structure. Group signature was generated by normalizing all features as a series of Z scores per condition (far right upper graph). Hierarchical clustering of conditions (y axis) and features (x axis) was performed using Partek and presented as heatmaps (far right lower graph). B. Illustration of the angle criteria used to define an interaction; angle subtended (α, β or γ) >0. C-D. Total number of encounters as a function of encounter duration in representative movie of socially raised WT flies (C), and when adding a 60-frames gap requirement between interactions (D). Black line represents the threshold (60 frames) under which encounters are not considered interactions for network analysis. E. Directed interactions quantified as the total duration between each pair of flies. F. Directed interactions quantified as the total number of interactions between each pair of flies.

We used the following requirements for an interaction: (1) Consistent with basic interaction criteria described by Schneider et al^70^ and based on the fact that 95% of social interactions (approach, touch and social clustering) occur in the range of 1-8 mm (fig. S1 A-C), we set the distance threshold for interaction between two flies to be 8mm or less, which is average of two body lengths. (2) the visual field of view of the focal fly is occupied by the other fly (angle subtended>0), indicating that the focal fly can see the other fly (Figure 1B). To minimize the number of false positives (random interactions), we required the angle and distance criteria be maintained for at least 2 seconds (Fig. 1C). This resulted in a large number of very short interactions, some of which could actually be long interactions that are mistakenly recognized as separate short interactions, due to small numbers of intermittent frames in which one of the conditions is not met (Fig. 1C). To resolve this, we added an additional requirement of a minimal time interval (gap) below which a subsequent interaction is considered an extension of the previous interaction between the same pair of flies. To find the optimal gap length, we tested a series of interaction and gap lengths and eventually selected a gap length of 4 s (120 frames) (Fig. S1D), which substantially reduced the number of very short interactions (Fig. 1D). We used weighted networks to account for the between-dyad variation in total interaction times over each test, and to avoid network saturation, an inherent limitation of binary networks. Next, we analyzed the symmetry level between interacting flies, by testing whether the total amount of time in which individual (X) interacts with individual (Y) correlates with the total amount of time in which individual (Y) interacts with individual (X). Performing this for all pairs of flies within each group resulted in high correlation (Fig. 1E), which was also apparent when quantifying total number of interactions between each pair (Fig. 1F). This suggests symmetric interactions over the course of the test, making directed analysis redundant in this setup. We used the interaction data to calculate 4 network features; Strength, Density, Betweenness Centrality and Modularity (Schematic illustration and explanation of the features are depicted in Figure 2I). In total, our data analysis pipeline generates 60 features that represent the behavioral repertoire of individuals within a group and their corresponding social networks. To process and analyze such rich datasets, we generated a comprehensive representation of all features using normalized Z-score scatter plots and hierarchical clustering to compare between experimental groups and highlight similarities and differences between them (Fig. 1A).

**Figure 2:**
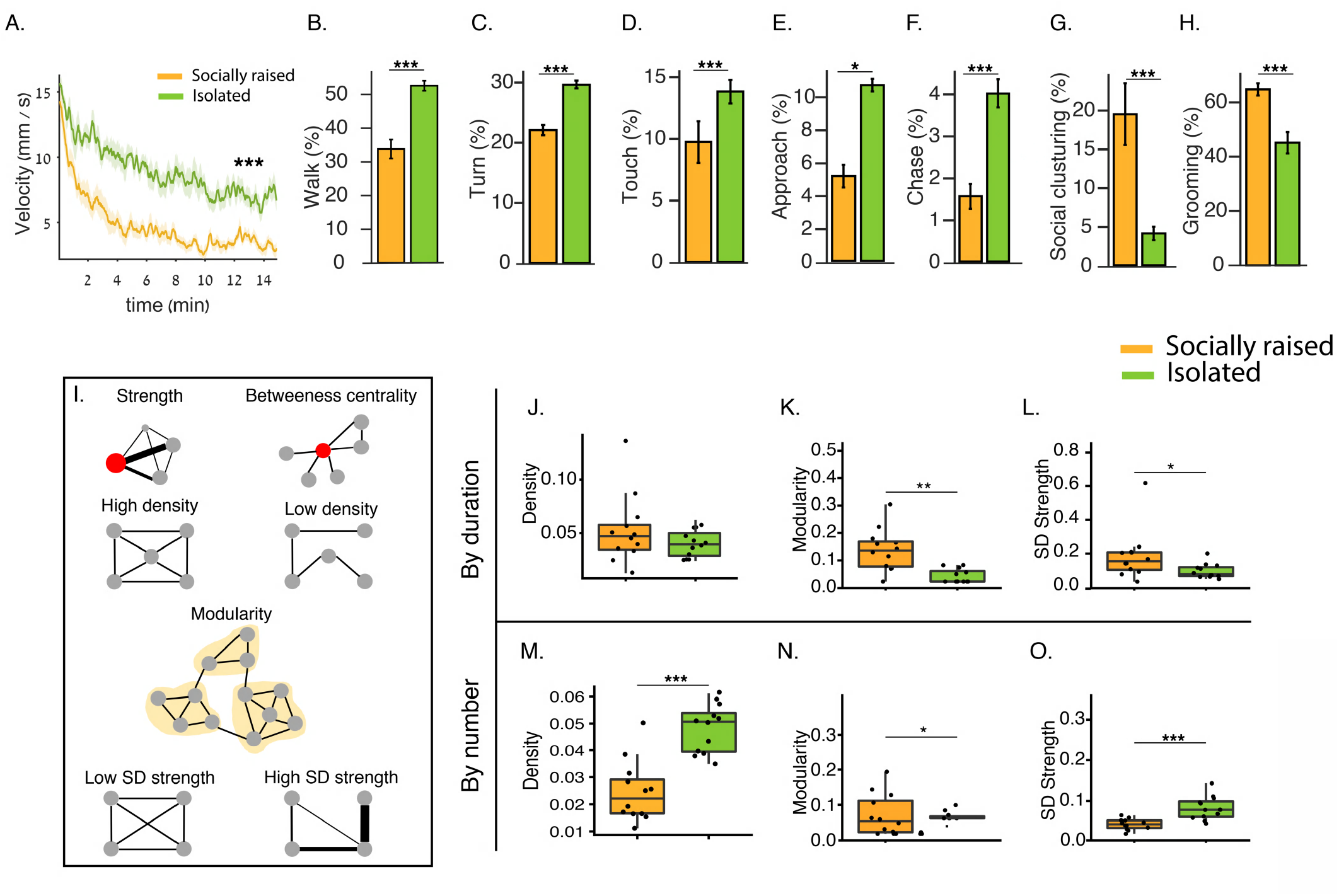
Prior social interaction in a group facilitates the formation of ordered social structures. A. Average velocity per-frame of previously isolated male flies (green) vs. socially raised male flies (orange) over 15 minutes. B-H. Average percentage of time previously isolated male flies (green) vs. socially raised male flies (orange) perform walk (B), turn (C), touch (D), approach (E), chase (F), social clustering (G) and grooming (H) behaviors. I. Illustration of network parameters; Strength is proportional to vertex size. Betweenness centrality is a measure of the tendency of the individual to serve as a hub connecting different sub-groups (high in red individual). Density of networks represents how saturated they are compared to the maximum possible. Modularity is a measure of the division of a network into sub-networks. Standard Deviation (SD) strength is a measure of the heterogeneity of the connections between individuals. J-O. Network density, modularity and SD strength calculated by network weights according to duration (J-L respectively) or number of interactions (M-O respectively) between previously isolated (green) and socially raised (orange) WT male flies. N=18, Wilcoxon test and FDR correction for multiple tests * P<0.05, ** P<0.01, *** P<0.001. Error bars signify SEM.

### Prior social interaction in a group facilitates the formation of ordered social structures

To test whether social experience can drive divergent forms of group behaviors, we generated two cohorts of wild-type (WT) Canton S male flies; one cohort of flies raised for 3 days with nine other flies (as groups of 10 male flies), while the other cohort raised in complete social isolation upon eclosion. After 3 days, 10 flies from each cohort were introduced into Fly Bowl arenas and their behavior was recorded for 15 minutes and analyzed (Fig. 1A). The two cohorts exhibited distinct repertoires of behavioral responses upon interaction with other flies in a group; socially raised flies displayed lower average activity levels, manifested by lower average velocity (Fig. 2A), shorter time spent walking (Fig. 2B) and fewer body turns than isolated male flies (Fig. 2C). Analysis of specific social behaviors revealed that socially raised flies exhibited less touch behavior (Fig. 2D), were less engaged in active approach (Fig. 2E) and spent less time chasing (Fig. 2F). Socially raised flies also spent more time grooming than isolated flies (Fig. 2H). Analysis of average duration (bout length) and frequency of specific behaviors revealed that touch, chase, approach, grooming and social clustering behaviors were significantly different between the two cohorts (Fig. 3A, Fig. S2A–H). Interestingly, average bout duration of approach behavior was similar between the two cohorts, while its frequency was higher in isolated flies (Fig. 3A and Fig. S2A, E), suggesting the difference in their social experience did not affect the duration of social encounters, but rather the frequency at which they occur.

**Figure 3.**
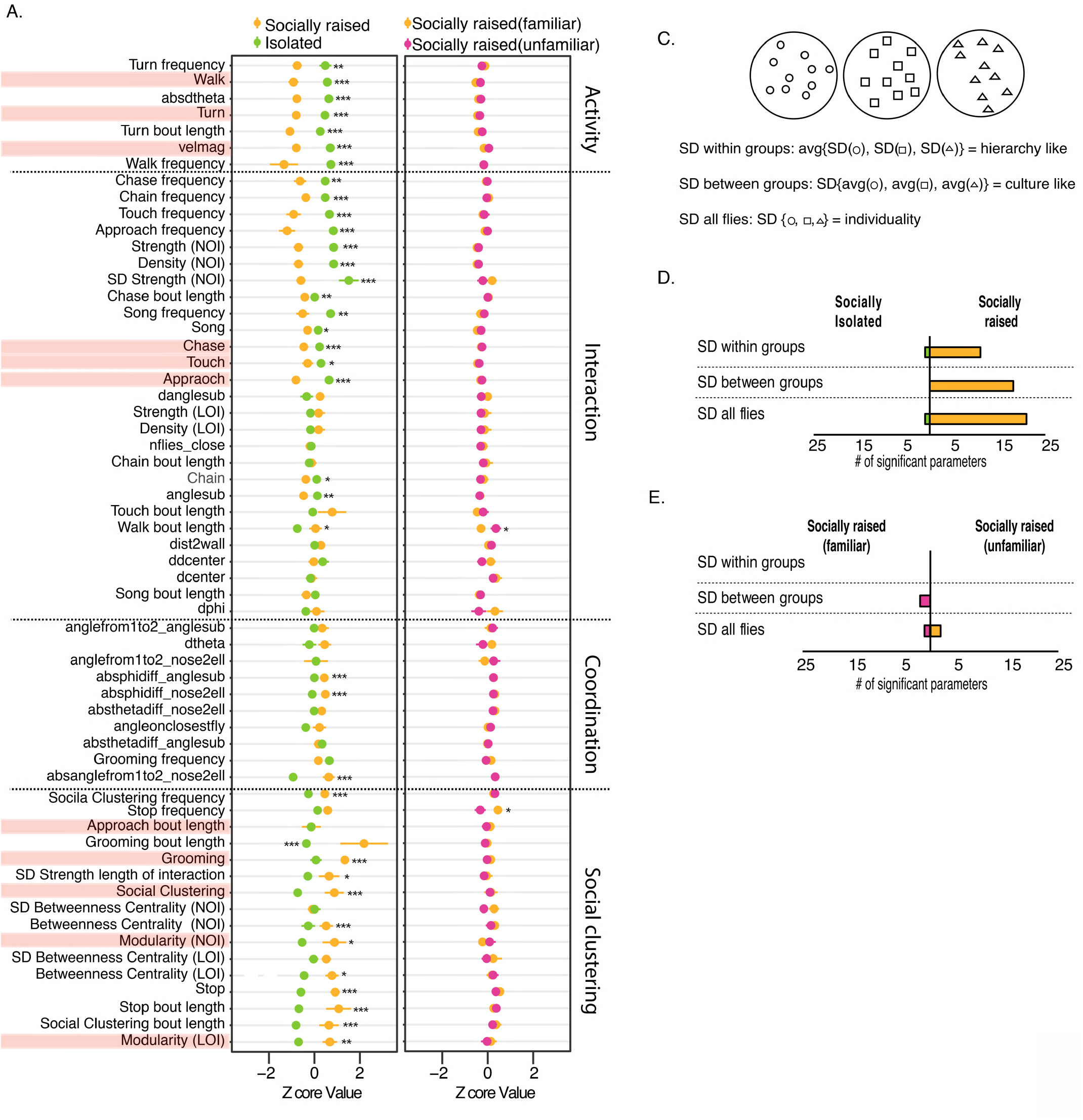
Social experience facilitates distinct group signature and increases behavioral variability. A-B. Behavioral signatures of previously isolated vs. socially raised WT male flies (A) and familiar vs. unfamiliar raised WT flies (B). Data is represented as normalized Z scores of 60 behavioral (A: N=18. B: N=25 t-test for normally distributed parameter or Wilcoxon test for non-normally distributed parameters. P-values were corrected using FDR. * P<0.05, ** P<0.01, *** P<0.001). Features mentioned in the results section are highlighted in pink. C. Graphical illustration of measuring variance within groups, between groups and across all individuals (all flies) in each condition. D-E. Number of behavioral features that display significantly higher variance and their SD is at least two-fold higher when comparing isolated to raised (D) and familiar vs unfamiliar (E). Statistical analysis was performed on SD of the entire population (all flies) (F test), SD of repetitions in each condition (between groups) (F test) and average SD within each repetition per condition (inside groups) (t-test). P-values were corrected using FDR. *P<0.05, **P<0.01, ***P<0.001. Error bars signify SEM.

The difference between socially raised and socially isolated flies can result from inherent differences in the kinetic properties of individuals, or from an emergent property of flies interacting in a group. To distinguish between these two possibilities, we compared the behavior of socially isolated and raised flies that were tested singly. If the differences between the groups stem from inherent differences in the kinetic properties of individuals, we would expect to identify kinetic differences between the two cohorts of singly tested flies. Remarkably, we did not observe any significant differences between the two cohorts, suggesting that the effects of social experience on behavior are an emergent group property expressed during group interaction (Fig. S3A-I). Another example for a difference in the emergent properties of socially raised and isolated groups is the tendency of socially raised flies to concentrate in certain zones within the arena, forming semi-stable social clusters consisting of three or more flies (Fig. 2G, Fig. S2M). This behavior was not apparent in male flies that raised in social isolation prior to testing, suggesting this behavior emerges from the social experience of flies rather than from the context of the behavioral test itself (Fig. 2G).

To investigate how group structure is affected by social history, we analyzed the network structures of groups composed of socially raised or socially isolated individuals. We calculated network weights according to the overall duration of interactions (emphasizing long-lasting interactions) or the overall number of interactions (emphasizing short interactions) between each pair of flies. Analysis by duration revealed that socially raised flies displayed higher modularity (Fig. 2K), SD strength (Fig. 2L) and betweenness centrality (Fig. S2L), suggesting that prior social experience promotes the formation of subgroups. Network analysis by number of interactions, which assigns equal values to long and short interactions and thus undervalues social clusters (Fig. 2J-L vs. M-O), revealed that the social networks of isolated flies are characterized by higher density (Fig. 2M), SD strength (Fig. 2O) and strength (Fig. 3A), suggestive of overall more interactions. In contrast, networks of socially raised flies have higher modularity (Fig. 2N) and betweenness centrality (Fig. 3A), similar to the results obtained with analysis by duration of interaction. Taken together, these differences indicate that socially isolated flies perform more short interactions compared to socially raised flies, while socially raised flies form networks with higher-order structures compared to those formed by isolated flies. Overall, these results show that the behavioral group signature of socially raised flies differs from that of previously isolated ones (Fig. 3A).

### Behavioral signature of socially raised flies does not require individual recognition

It is plausible that the observed differences between socially raised and isolated cohorts result from the familiarity of raised flies with the individuals they are tested with. Therefore, we asked whether the distinct features exhibited by socially raised males result from their familiarity with individual members that occurred during housing, or from the internal state associated with the general experience of living in a group. To distinguish between these two possibilities, we tested socially raised flies with either familiar or unfamiliar individuals. One cohort was tested with the same flies they were previously housed with (familiar), while the other cohort was tested with socially raised flies from other groups (unfamiliar). Encountering familiar or unfamiliar flies did not result in different behavioral signatures (Fig. 3B), suggesting that the dynamics captured during the test result from the general experience of interacting with others rather than by specific previous interactions. We next tested whether other conditions that are known to modulate internal state such as repeated ethanol exposure, starvation, and different circadian time shifts, also affect group interaction. We did not observe any significant difference between these conditions and their controls (Fig. S5), implying that not all experiences that modulate internal state affect group dynamics in the context used in our experimental paradigm.

### Prior social interaction increases behavioral variability

The existence of a complex social structure in groups of socially raised flies suggests that in addition to the observed differences in the means of various behaviors, there may be additional effects on the distribution of certain features. Indeed, when analyzing the behavioral signatures of socially raised and isolated male flies, we observed that socially raised flies exhibited higher variance across several behavioral features (Fig. 2, 3A; compare error bars). To further investigate this, we compared the variance of all behavioral features between groups of socially raised and isolated male flies. We analyzed the variance of each behavioral feature in three ways: (a) average standard deviation (SD) of each group (each movie), reflecting variation inside each group (SD within groups, Fig. 3C); (b) SD of averages between experimental groups per condition, reflecting variation between groups (SD between groups, Fig. 3C); and (c) SD across all flies per condition, reflecting individual differences between all flies regardless of groups (SD all flies, Fig. 3C). We documented a higher number of behavioral features that displayed significantly higher variance (SD two-fold higher in one condition + statistically significant) in socially raised flies between groups (18 out of 56 parameters; Fig. 3D), within groups (11 out of 56 parameters; Fig. 3D) as well as between all flies (21 out of 56 parameters; Fig. 3D). This indicates that the behavior of socially raised flies is more diverse than that of isolated flies, possibly reflecting a broader repertoire of behaviors in individuals which is shaped by prior interactions during the experience phase. Increased variability between groups of socially raised males that have presumably had identical experience suggests that each group possesses distinct group characteristics that were shaped during the housing period before the test. To test this hypothesis, we asked whether between-group variance stems from inter-individual recognition or is based on the general experience of living in a group. For that, we performed a similar analysis in male flies that were housed in groups and tested either with the same group members or with flies that were housed in other groups (data taken from the experiment of Fig. 3B). We documented very few parameters that were distributed differently between flies tested with familiar or unfamiliar flies, implying that the general experience of living in a group also shapes the variance of behavioral responses, and that individual recognition has little to no effect on behavioral variance in a group (Fig. 3E).

### Visual cues are necessary for expressing the behavioral signature of socially raised flies

So far, we have shown that different types of social history can form divergent group dynamics and structure. Next, we set out to dissect the sensory elements required for the expression of such differences. We started by assessing the role of visual cues in forming specific behavioral signatures during the test. For that, we analyzed the behavior of socially raised flies in light or dark conditions (this did not interfere with tracking since recording is performed using IR backlight). Socially raised flies that were tested in the dark displayed more walk, turn and touch behaviors than those tested in the light (Fig. 4A), and spent a larger fraction of time in chase and approach behaviors, while showing less social clustering and grooming behaviors (Fig. 4A). Moreover, approach behavior in the dark was significantly longer and more frequent than that in the light (Fig. 4A), while frequency and duration of social clustering was lower in the dark. Interestingly, although the average velocity of flies in the presence or absence of light was not statistically different (Fig. 4A), flies tested in the light reduced their velocity over time, while flies tested in the dark maintained a constant velocity for the duration of the experiment. This was also evident in several other behavioral features, such as walk and turn behaviors, suggesting that flies habituate to environmental conditions in the light but not in the dark (Fig. S6A-F). Network analysis revealed lower SD strength and betweenness centrality in groups tested in the dark, by analysis of duration of interactions (Fig. 4A), while analysis by number of interactions revealed that flies in the dark display higher density, strength and SD strength than flies in the light (Fig. 4A). Therefore, we postulate that light is required for the group signature of socially raised male flies.

**Figure 4:**
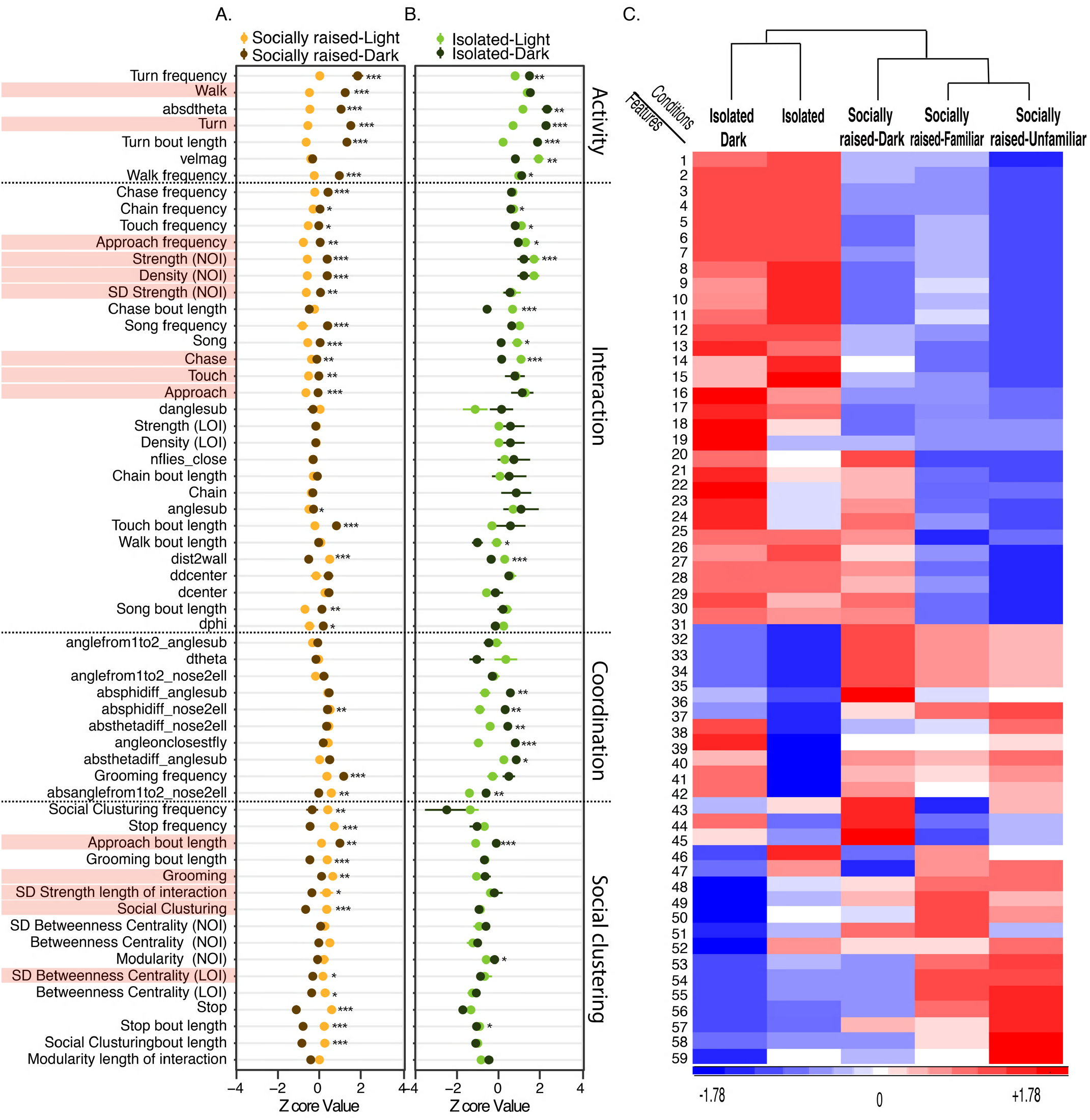
Visual cues are necessary for expressing the behavioral signature of socially raised flies. A-B. Behavioral group signatures (represented as normalized z-scores) of socially raised (A) or previously isolated (B) WT male flies tested in normal lighting conditions (light) vs. light deprivation (dark). LOI - calculated according to the length of interactions. NOI - calculated according to the number of interactions. N=18 and 10, respectively. t-test for normally distributed parameters or Wilcoxon test for non-normally distributed parameters. P-values were corrected using FDR. *P<0.05, **P<0.01, ***P<0.001. Error bars signify SEM. Features mentioned in the results section are highlighted in pink. C. Hierarchical clustering (dendrogram) of group signatures of the following experimental conditions: socially raised (raised familiar), unfamiliar socially raised (raised unfamiliar), socially raised tested in dark (raised dark), socially isolated tested in light (isolated) and socially isolated tested in dark (isolated dark). List of numbers represent behavioral features. Full list in Fig. S4B.

We next aimed to uncouple the behavioral changes observed during light deprivation: those that result from the role of visual cues in a typical social interaction in a group, from those that specifically depend on prior social experience. For that, we tested groups of socially raised and socially isolated flies in the presence or absence of light (Fig. 4A, B). Behavioral features that are affected equally by light in both groups, represent features that are light-dependent but not sensitive to social experience, while features that are only affected in one group are those that turn into light-dependent by previous social experience. To visualize this, we plotted distinct features that are influenced by visual cues in each condition. We identified 22 unique features that are sensitive to visual cues only in socially raised flies, and only seven in isolated flies, suggesting that the experience of an enriched social environment requires light to be fully expressed (Fig. S4A). Most features unique to the socially raised group are associated with social clustering (reduced in the absence of light) and interaction (increased in the absence of light). The opposite regulation of these two types of features suggests that in the absence of light, socially raised flies undergo a shift from a quiescent state to a more active state, characterized by more approach, chase and touch behaviors. In contrast, groups of previously isolated flies displayed a decrease in a few interaction-related parameters and an increase in a class of parameters that reflect changes in angle and speed between two close individuals in the absence of light (absanglefrom1to2, absphidiff, absthetadiff and angleonclosestfly; see Table S1 for more details) (Fig. S4A). This may signify an increase in coordination between pairs of flies and suggest that isolated flies in the dark generally tend to be less mobile but more engaged with others when interacting (Fig. 4B, Fig. S4A).

To assess whether the group signatures of these conditions reveal an underlying similarity, we performed hierarchical clustering analysis on group signatures of all conditions (Fig. 4C, list of features in Fig. S4B). This analysis revealed two main clusters based on social history; one of conditions in which flies were isolated prior to the test and another of conditions in which flies were socially raised. Interestingly, socially raised flies that were tested in the dark did not cluster with either groups, reinforcing the notion that specific visual cues are necessary for the expression of group signatures associated with social experience, but are not sufficient to fully shift group signature from that of socially raised to that of isolated.

### cVA perception via Or65a neurons shapes social group interaction

In addition to visual cues, a central element in social interaction of flies is pheromone-based communication. The male-specific pheromone cVA is known to mediate experience-dependent changes in aggressive behavior, where chronic exposure to cVA found on conspecifics during group housing, is known to reduce male–male aggression^61,81^. cVA is perceived via two olfactory receptor neurons (ORNs): Or67d, which mediates acute responses to cVA, and Or65a, which mediates chronic responses to cVA^61,82^. We investigated whether cVA perception impacts the group signature of socially raised flies. For that, we blocked cVA perception by constitutively expressing the inward rectifying potassium channel Kir2.1 in Or65a- or Or67d-expressing neurons of socially raised flies and then analyzed their group behavior. Inhibition of Or67d neurons did not lead to major differences between experimental flies and genetic controls, suggesting that the function of Or67d neurons is not necessary for the formation of the behavioral signature associated with social group experience (Fig. 5A). In contrast, inhibition of Or65a neurons dramatically changed the group signature of socially raised flies, increasing average velocity and overall time flies engaged in approach, chase and touch behaviors compared to genetic controls (Fig. 5B). Network analysis revealed higher strength and lower betweenness centrality in the Or65a experimental group compared to genetic controls, by both duration and number of interactions (Fig. 5B). Overall, this suggests that Or65a-but not Or67d-expressing neurons function in shaping the group behavior of socially raised flies.

**Figure 5.**
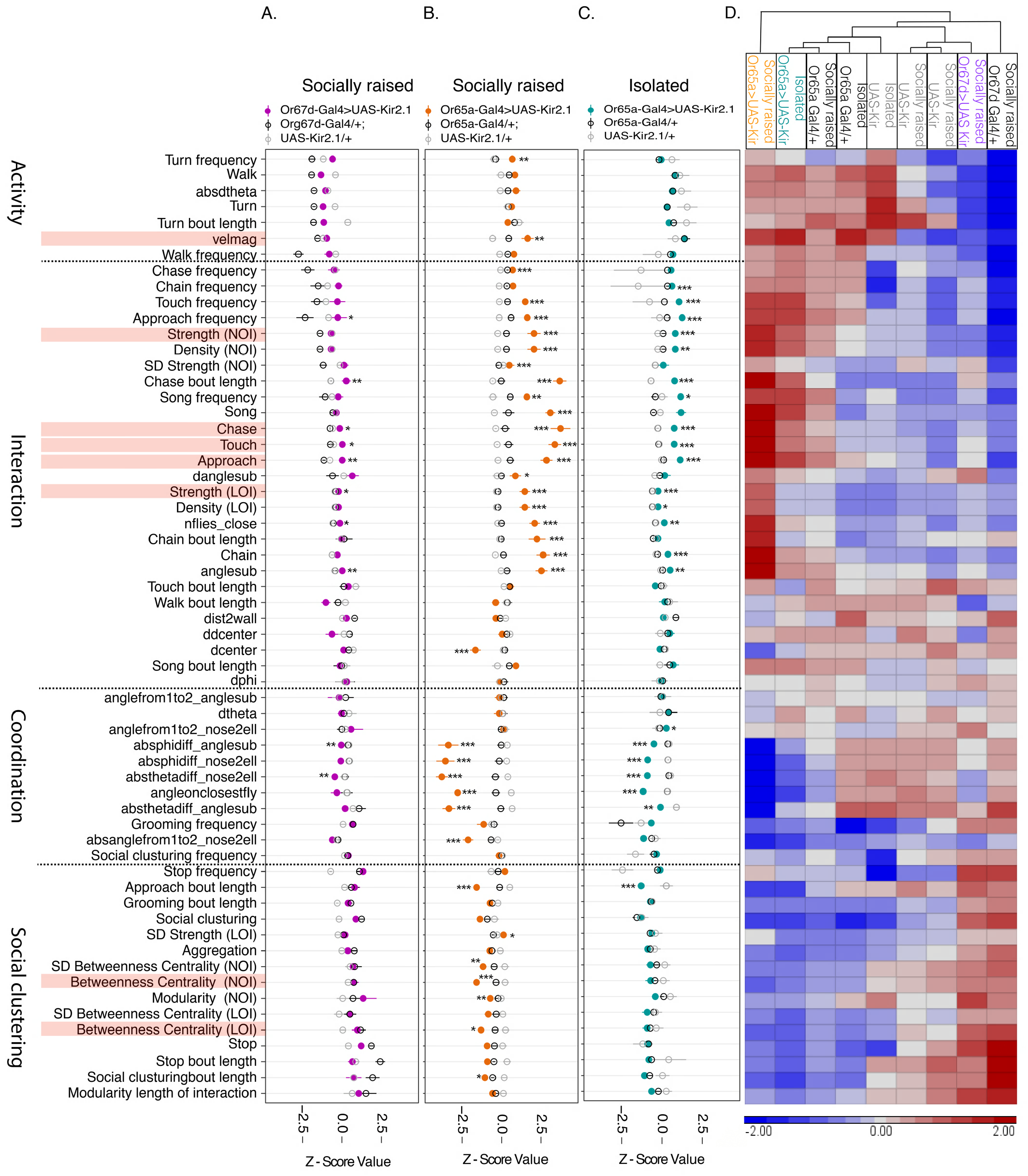
cVA sensing via Or65a neurons shapes social group interaction. A-C. Behavioral group signatures (as normalized z scores) of socially raised Or67d-Gal4/+; UAS-Kir2.1/+ flies compared to genetic controls (A), of socially raised Or65a-Gal4/+; UAS-Kir2.1/+ flies compared to genetic controls (B) and of previously isolated Or65a-Gal4/+; UAS-Kir2.1/+ flies compared to genetic controls (C). LOI - calculated according to the length of interactions. NOI - calculated according to the number of interactions. N=7, 13, and 8 respectively. One-way ANOVA with Tukey’s range test for normally distributed features or Kruskal Wallis followed by Wilcoxon signed-rank test for non-normally distributed features. P-values were corrected using FDR. *P<0.05, **P<0.01, ***P<0.001. Error bars signify SEM. Features mentioned in the results section are highlighted in pink. D. Hierarchical clustering (dendrogram) of behavioral group signatures of all experimental conditions in A-C.

This experimental design does not distinguish between the role of Or65a neurons during experience and test phases, due to the constitutive nature of this neuronal inhibition. To test the role of Or65a neurons during the test phase, we performed a similar experiment in isolated male flies, which are expected to be exposed to cVA only during the test. If Or65a expressing neurons function only to shape the group signature of socially raised flies via exposure to cVA during the experience phase and before test, we expect the inhibition of these neurons not to affect the behavioral signature of isolated flies. Surprisingly, inhibition of Or65a neurons in isolated male flies resulted in changes of several behavioral features, although Or65a neurons are thought to only mediate chronic responses to cVA over long time courses^61^. Experimental flies (*Or65a>Kir*) exhibited more touch, approach, chase and chain behaviors than genetic controls, and increased network strength as measured by duration of interaction (Fig. 5C). However, these effects were less extreme than those displayed by socially raised male flies (Fig. 5B vs. 5C). This unexpected result suggests that Or65a neurons mediate acute as well as chronic responses to cVA.

Interestingly, some effects of Or65a neuronal inhibition are identical between socially isolated and socially raised flies, including a decrease in three coordination-related parameters (Fig. S7A–C) and a significant increase in chain, chase, chase bout length, touch and approach behaviors (Fig. S7D–H). Moreover, both experimental groups displayed higher network strength (measured by duration of interaction, Fig. S7I), suggesting that inhibition of Or65a neuronal activity facilitates behaviors that are associated with social isolation. Overall, although these two conditions share similarities, the effect of Or65a inhibition was more profound in socially raised flies than in socially isolated flies, reflected by the higher number of behavioral features affected (35 vs. 22 out of 60, Fig. 5B, C). Hierarchical clustering analysis between conditions revealed that flies in which Or67d neurons were inhibited are similar to their corresponding genetic controls, supporting the conclusion that Or67d neurons do not mediate behavioral responses of socially raised male flies in a group (Fig. 5D). In contrast, socially raised male flies in which Or65a neurons were inhibited are clustered apart from their genetic controls and all other tested conditions, indicating that cVA perception though Or65a sensing neurons is necessary for the formation of a certain internal motivational state via the experience of group housing, leading to a specific group signature (Fig. 5D).

### Heterogenous groups of flies exhibit dynamic social interaction that is shaped by group composition

So far, we have used homogenous groups of flies which were subjected to environmental or genetic manipulation as a tool to investigate the interplay between social experience and the resulting group behavior. This approach eliminates the inherent contribution of inter-individual differences to group structure, which proved valuable in dissecting the elements that shape social group behavior. Next, we asked how the dynamics inside the group are shaped by different individuals. For this, we generated groups that contain varying ratios of male flies with two distinct states: socially raised flies and hyper-aggressive isolated flies. Hyper-aggressive male flies were generated by knocking down (k.d) *Cyp6a20* (a manipulation known to induce aggression)^20^, and keeping these flies isolated from eclosion. We postulated that highly aggressive k.d flies would disrupt collective-like group behaviors such as social clustering and thus change the behavioral signature of the group.

To verify that these flies indeed behave as expected, we tested their social interaction in groups of flies, and compared it to *Cyp6a20* k.d flies that were socially raised before the test and to that of socially raised WT control flies (Fig. S8). We did not document any difference between the two cohorts of *Cyp6a20* k.d flies. However, compared to the WT control group, both *Cyp6a20* k.d cohorts displayed more walk, turn and chase behaviors (Fig. S8 B,C,F), while exhibiting lower social clustering and grooming behaviors, as expected (Fig. S8 G,H). This suggests that the genetic manipulation in this case eliminates the effects of previous social experience on group signature.

Next, we introduced increasing numbers of hyper-aggressive flies into groups of socially raised WT male flies (10%–50% of the total number of individuals) and measured their group behavior. The behavior of each experimental group was normalized to a control group of 100% socially raised WT flies which was tested at the same time, enabling statistical comparison of all behavioral features across all experimental groups (0-50%), unlike previous experiments in this work which can only be compared to their controls. To gain a general overview of the patterns associated with gradual changes in group composition, we examined the normalized behavioral signatures using hierarchal clustering (Fig. 6A). Overall, the conditions are clustered into two main branches: one containing the homogenous WT group with the 10%–30% mixed ratio groups, and a separate branch containing groups of 40%– 50% mixed ratios, suggesting a behavioral transition from homogenous to 50% mixed ratio groups. The differences between these two extremes resemble those of socially raised vs. socially isolated flies, suggesting that the addition of 50% aggressive flies is sufficient to convert group behavior into that of a social isolation-like state (Fig. 4C vs. Fig. 6A). Overall, clustering of features suggests a somewhat gradual transition from 0 to 50%. This trend is best demonstrated by the increase in the number of features that exhibit a significant difference compared to 100% WT flies (Fig. 6B). We identified a suit of features associated with an increasing number of *Cyp6a20*-knockdown (KD) flies: a cluster of decreasing features and a cluster of increasing features (Fig. 6A). Some decreasing features corresponded to social clustering and network structure, while increasing features were related to activity and interaction (Fig. 6A). Some of these features exhibited a gradual change as the number of *Cyp6a20*-KD flies in a group increased. These included a gradual decrease in social clustering, grooming, stop, and stop bout length (Fig. S9A–D), and a gradual increase in walk, angular speed (absdtheta), turn, and turn bout length (Fig. S9E–H). Interestingly, some behavioral features showed parabolic-like changes across increasing ratios of *Cyp6a20*-KD flies, with maximal or minimal values at 20%–30%, including touch behavior and several other features expected to be associated with coordination between two individuals (absphidiff_nose2ell, absphidiff_anglesub; Table 1). Some behavioral features were more sensitive than others to changes in group composition, such as grooming, approach and turn behaviors, which were significantly different from controls even at 20% mixed ratio, while other features such as social clustering exhibit a significant change only at 40-50%. This suggests that changes in the level of approach behavior within a group precede changes in more collective-like behaviors such as social clustering (Fig. 6A).

**Figure 6.**
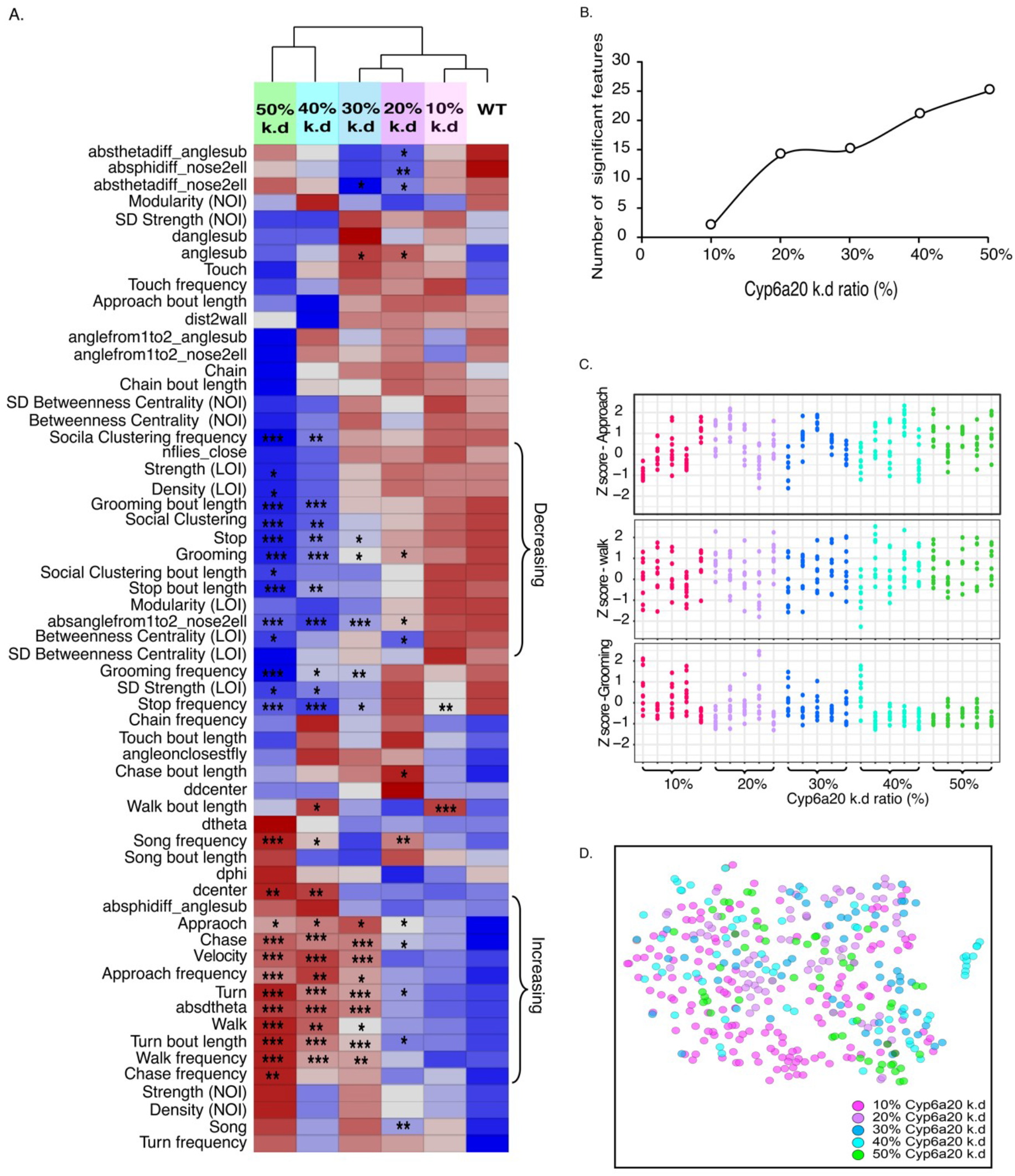
Sub populations of aggressive flies in a group affect different features of group behavior. A. Hierarchical clustering of behavioral signatures of groups composed of different ratios of socially isolated Cyp6a20 -Gal4/+; UAS-Cyp6a20-RNAi/+ and socially raised WT flies (0-50%). LOI - calculated according to the length of interactions. NOI - calculated according to the number of interactions. Data of each experimental group was normalized to a WT control group which was tested at the same time. To compare log-ratios of means (test/control), all values were log2-transformed and statistically tested as mean log-values. The effect of treatment and mutant number on the fraction of each parameter was tested with a linear regression and a 2-way ANOVA was performed on the resulting model. Log-ratios between different number of mutants were compared in terms of difference of differences defined with by linear contrasts, FDR correction was applied to all comparisons. * P<0.05, ** P< 0.01, *** P<0.001 N=14, 8, and 6 for groups of 10%, 20-30% and 40-50%, respectively. B. Number of significantly different behavioral features compared to controls as a function of the ratio of isolated *Cyp6a20* k.d - to raised WT flies in a group (10-50%). C. Per-fly distribution of three normalized behavioral features (interaction, walk, grooming) in groups containing increasing ratios (0-50%) of isolated *Cyp6a20* k.d to socially raised WT flies. Each column represents individuals as dots in one movie. Analysis of the distribution inside each group is not significantly different between conditions (F test, n.s.) D. t-SNE analysis of all individuals in 10-50% groups across all behavioral features.

It could be argued that the behavioral pattern exhibited by mixed groups represents an average of two distinct subgroups and not an integrated structure of all individuals within the group. If so, the differences observed at the group level would result from the existence of *Cyp6a20*-KD flies having higher values of approach behavior and lower values of social clustering, which would drastically affect the group average, depending on their relative ratio within the group. To test this, we analyzed the per-fly distribution of each condition. If each group is composed of two distinct subgroups (WT and *Cyp6a20*-KD flies), we would expect this to be reflected in a bi-modal distribution, which would become more pronounced as the ratio of *Cyp6a20* k.d flies increases. Single-fly analysis of features that exhibit changes with an increased number of mutant flies, such as walk, approach and social clustering, did not show a bi-modal distribution, making it impossible to identify subgroups that correspond to mutant or WT flies (Fig. 6C, f-test). To further analyze the distribution of group members in these mixed-ratio groups, we use t-SNE, a dimensionality reduction technique, to analyze all individuals across all features, which failed to depict any clear existence of subgroups (Fig. 6D). This finding suggests that both WT and mutant flies change their behavioral responses when interacting in a group to generate a single behavioral signature, implying that group structure and dynamics reflect a level of complexity that cannot be explained as a simple average of the individuals that constitute it.

## Discussion

Understanding the principles underlying the complex nature of social group interaction is conceptually and computationally challenging. In this work, we simplified this complex phenomenon to a series of experiments in which we controlled the social experience and internal states of individuals within a group to illuminate patterns representing distinct structures and behavioral responses of groups under different social conditions. Each condition was represented by a “group signature” containing a collection of 60 distinct social network and behavioral features. This comprehensive analysis provided a broad examination of behavioral states, highlighting similarities and differences between groups, confirming our initial hypothesis that different social histories give rise to the formation of distinct and robust group signatures, that are indicative of specific social group structures. We showed that groups composed of socially raised male flies exhibit social clusters and high network modularity, indicating the existence of stable subgroups and ordered social structure that are not apparent in groups of isolated flies. Some of the observed differences between the groups of socially raised and socially isolated flies satisfied our initial predictions, such as the increased activity in isolated flies and increased social interaction, as well as the formation of social clusters in the socially raised group due to reduced aggression. On the other hand, the prediction that isolated flies will exhibit social avoidance was not supported. In fact, socially isolated flies displayed higher number of interactions, approaches, and network density.

Using hierarchical clustering to compare between group signatures allowed us to identify specific elements which are shared across conditions. For instance, clustering of socially raised flies tested in the dark with that of previously isolated flies highlights the contribution of visual cues to the expression of group signatures, whereas clustering analysis of flies in which cVA sensing neurons were inhibited suggests that cVA perception shapes group structure during experience phase and during test. Moreover, the analysis of group signatures revealed two aspects relevant to the connection between sensory information and behavior: (a) existence of behavioral features that are “primed” by social experience to become light-dependent (i.e. social experience affects their light-dependence); (b) an emerging role for *Or65a* expressing neurons in regulating acute male–male interactions in addition to its well-established role in suppressing aggression upon long exposure to cVA^61^ or possibly a cVA independent role. Accordingly, hierarchical clustering indicated that inhibition of Or65a neurons affected many features in socially raised flies, some of which were also changed in isolated flies and are associated with increased activity in both cohorts. These common features are higher in isolated experimental flies when compared to their corresponding genetic controls, suggesting a role for Or65a neurons in reducing activity levels during the test.

Based on evidence suggesting that inter-individual recognition plays a role in male-male aggression encounters^83^, we expected recognition to shape also social interaction in of flies. We found no evidence for a role of inter-individual recognition in the formation of groups composed from socially raised flies, suggesting that although recognition is valuable in the context of aggression over limited resources, the context use d in our study is not sufficient to measure its importance. This finding is consistent with studies in social insects demonstrating that collective group behaviors do not require individual recognition^5^. Another example for the role of context to the expression of behavior is seen in the emergent differences in group behavior between groups of socially raised and isolated flies that are only evident in group context and not when the flies are tested alone. This fits well the conceptual model proposed by Anderson and Adolphs for the interplay between emotional behaviors and distinct internal states^11^, suggesting that group signatures integrate the expression of internal states, shaped by experience, with the specific context in which group behavior is measured.

The differences in variance between socially raised and isolated flies indicate that early-life experiences can modulate behavioral variability within and between groups. Inter-individual variability is a broad phenomenon documented in many species^84–92^, and was shown recently to be under neuromodulation in *C. elegans*, suggesting that behavioral variability is a biologically regulated process^93^. The functional importance of such variability can be seen in *Drosophila* studies demonstrating that increased behavioral variability can contribute to fitness^94^. Notably, our results also reveal increased variability between groups of socially raised flies, suggesting that social experience increases the repertoire of possible group phenotypes, the functional outcome of which remains to be studied.

Using network analysis as a tool to quantify social structures, we show that certain aspects of group structure are modulated by the social history of individuals that compose the group. Previous studies in *Drosophila* used social network analysis to dissect the principles that shape social interaction^13,70^. Interestingly, although the presence of visual cues affected several network features in our behavioral setup, Schneider et al. reported no effects of the absence of light on network structure^70^. This apparent discrepancy between our study and that of Schneider et al. could result from different approaches when measuring network structure (binary vs. weighted); while both studies documented shorter interactions in the absence of light, the effect on network structure is only evident when using weighted networks.

Studies of collective behaviors in various animals including honeybees, ants, birds and fish exemplify synchronization as a key component of collective behavior^1,5,95^. Although *Drosophila* do not display such a degree of collective/coordinated behaviors as these organisms, they do exhibit behavioral responses that involve collective features, such as different responses to threat when in a group, changes in memory retrieval that depend on social experience, cooperation in feeding behavior and even aggregation, suggesting the existence of a collective response that can increase survival or reproductive success^4,55,72,96–102^. Adding to this, our results demonstrate the presence of social clusters, characterized with increased coordination between individuals, stable distances between individuals, long-lasting interactions, which are correlated with increased grooming, all of which are suggestive of a semi-collective state, in agreement with previous studies^103,104^. We show that the degree of this highly social state strongly depends on prior social experience, and its expression requires cVA perception and visual cues. The existence of such an ancient form of coordinated behavior may serve to explore the neuronal and genetic mechanisms underlying collective behaviors, as suggested by de Bono^105^.

Lastly, we demonstrate that group behavior and its corresponding structure depends on its composition. Hierarchical clustering of groups composed of different ratios of super-aggressive flies identified a cluster of features that is highly sensitive to changes in group composition. This cluster contains features associated with coordination between individuals and features associated with social clustering, implying that specific clusters of behavioral parameters within a behavioral signature may reflect changes in the ability of the group to form semi -collective structures^1^. Importantly, although the groups of mixed populations consist of two types of individuals that form distinct signatures when tested separately, their combination does not result in two distinct populations but rather a single close to normal distribution of all individuals within the group, as supported by^106^. This raises questions about the interactions and mechanism that facilitate the formation of unimodal distribution in groups composed of individuals with highly different internal properties.

The finding of state-dependent group signatures hints at the existence of distinct and consistent behavioral responses of groups to specific social conditions, which give rise to distinct group structures. These structures and their dependency on specific sensory information raise questions about the kinetics of their formation and the neuronal mechanisms that shape interactions that sustain such structures. These complex multi-sensory requirements also raise general questions about the ability of semi-natural social interactions such as technology based social communication platforms to fully mimic the complex repertoire of experiences associated with face-to-face interaction, as a prerequisite for the full expression of social group interactions.

## Acknowledgments

We thank all members of the Shohat-Ophir and Ilani labs for fruitful discussions and technical support. We specially thank Kristin Branson and Alice Robie (HHMI Janelia research campus) for their valuable advice and guidance with the experimental and computational design and for Karla Kaun (Brwon University) for her productive suggestions. This work was supported by the Israel Science Foundation Grant 384/14, Israel Science Foundation Grant 174/19 and Israel Science Foundation Grant 384/14.

## Author contributions

Conceptualization, A.B., S.B.-S., A.I. and G.S.-O.; Methodology, A.B., and S.B.-S. and E.C; Investigation, Software, S.B.-S.; Writing, A.B, A.I. and G.S.-O.; Statistical Analysis, J.B., Funding Acquisition, G.S.-O and A.I; Supervision, G.S.-O and A.I.

## Methods

### Tracking

Flies where inserted in groups of 10 into Fly Bowl arenas^107^, and 15 minutes of video was acquired with Fly Bowl Data Capture (FBDC)^79^ and analyzed using CTRAX^80^ to obtain flies’ orientation, position, and trajectories.

### FixTRAX

We programmed this additional software in MATLAB in order to fix CTRAX tracking errors. FixTRAX uses a set of assumptions to fix CTRAX output based on 4 types of errors we observed in our CTRAX output data, which mostly happen when flies are relatively immobile for long time periods and require correction prior to further analysis. The errors are: (a) unifying two or more identities when flies are close, (b) mistakenly identifying a dark spot as a fly, (c) not recognizing a fly for several frames and (d) not maintaining the same identities over the entire movie. FixTRAX uses two fix algorithms; a main algorithm and a subsidiary control algorithm (Supp FixTRAX code and user instructions). The main algorithm is based on finding a sequence of incorrect frames that represent one mistake, then creating a table from that sequence with statistical scores for every pair of identities: one that disappeared and another that appeared. This score represents the probability that the two identities represent the same fly. Based on their score, the algorithm decides which identities to unify and which identities are false and can be deleted. After unifying two identities, data for missing frames is computed according to the fly’s approximate location, calculated as the shortest path between start and end positions of that specific error. The subsidiary algorithm unifies each identity that disappeared with the first identity that appeared. Both algorithms stop when all identities are unified, and the number of identities matches the number of flies the user stated are in the video. FixTRAX selects the fix algorithm that was able to maintain the identities of all flies in the movie with minimal insertions or deletions of identities to the original tracking file. Finally, FixTRAX plots a graph of the number of identities that were added and deleted for per frame, which can help the user adjust CTRAX’s tracking parameters and the fix algorithm parameters to minimize tracking errors. Experiments which were not tracked correctly were discarded. Finally, FixTRAX output is converted into JAABA compatible output using the algorithm specified in Kabra et al.^78^ to generate general statistical features as in^80^ (Fig. 3A). FixTRAX error rate is presented in FixTRAX error rate supplementary file.

### Kinetic analysis

Scripts were written in MATLAB to use the JAABA code to generate the statistical features as specified in Kabra et al.^78^. Time series graphs (per frame) were created using JAABA Plot^78^.

### Quantification of specific behaviors

JAABA Classifiers^78^ were trained on various movies to identify specific behaviors: Walk, Stop, Turn, Approach, Touch, Chase, Chain, Song, Social Clustering and Grooming. Bar graphs were created using JAABA Plot^78^.

### Network analysis

An Interaction matrix was created in MATLAB (using the interaction parameters stated below) and saved as a text file. Two interaction matrices were created for each movie, one with the total number of frames each pair of flies were interacting divided by the number of frames in the movie and another with the number of separate interactions between each pair of flies divided by the maximum number of possible interactions, calculated as:

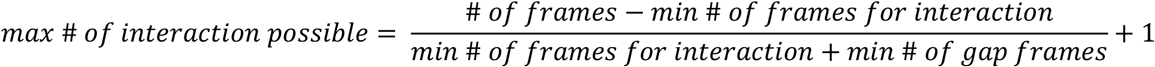

The parameters to define an interaction are: angle subtended by the other fly > 0, distance between the nose of current fly to any point on the other fly ≤ 8 mm, number of frames for interaction ≥ 60 and number of gap frames ≥ 120. Interaction end is defined when distance or angle conditions are not maintained for 4 seconds.

Networks and their features were generated from the interaction matrix in R using the igraph package^108^. The function that was used to the generate networks is “graph_from_adjacency_matrix” with parameters “mode = undirected” and “weighted = TRUE”. Density was calculated on all movies with the formula:

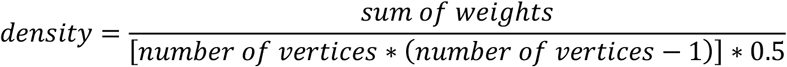

Modularity was calculated using the “modularity” function on output from the “cluster_walktrap” function^109^. Strength was calculated using “strength” function and SD Strength was calculated on all movies using “sd” function on the strength value. Betweenness Centrality was calculated on all flies using the “betweenness” function and SD Betweenness Centrality was calculated on all movies using “sd” function on the Betweenness Centrality value. Box plots were created using R.

### Variance analysis

Standard deviation (SD) of all flies was calculated as standard deviation of all per-fly data (all experimental repetitions together) for each feature per condition. SD between groups was calculated as standard deviation of all per-movie (experimental repetitions) averages for each feature per condition. SD within groups was calculated as the average of all per-movie standard deviations (variance within each experimental repetition) for each feature in each condition.

### Standardization and normalization

For all experiments except those of ratios of sub populations (Fig. 6), each feature was standardized according to all values calculated in our experiments for that feature to generate a z-score, as was done by Schneider et al.^70^. Scatter plots were created using R.

Sub populations experiment (Fig. 6): Each feature in every experimental group was first normalized to a control condition of 10 WT flies. Features were then standardized according to all normalized values of all other experimental groups to generate z-scores.

### Hierarchical clustering

Hierarchical clustering and heatmaps were created using Partek® software (Copyright, Partek Inc. Partek and all other Partek Inc. product or service names are registered trademarks or trademarks of Partek Inc., St. Louis, MO, USA). Each condition (heatmaps y axis) represents average standardized values of all repetitions.

### Fly lines

Flies were raised at 25°C in a 12-h light/12-h dark cycle in 60% relative humidity and maintained on cornmeal, yeast, molasses, and agar medium. Canton S flies were used as the wild-type strain. All transgenic fly lines were backcrossed at least 5 generations into a white Canton S background. Or67d-GAL4, Or65a-GAL4 and UAS-Kir2.1 fly lines were obtained from HHMI Janelia Research Campus. Cyp6a20-GAL4 was obtained from the Heberlein GAL-4 collection and Cyp6a20-RNAi was obtained from VDRC.

### Behavioral setup

Socially raised vs. Isolated: flies were lightly anesthetized with CO2 and collected shortly after hatching. Flies were then inserted into food vials, either alone (isolated) or as a group of 10 (raised) for 3 days, in a light/dark cycle of 12/12. The isolated flies were inserted into a food vial in a group of 10 and then loaded into the test arenas, same as experienced flies. All flies experienced similar habituation to the arena of about 1 minute.

Light vs dark: flies were collected as before and housed in groups of 10 or in isolation as before. During the behavioral test, light was off (dark) or on (light).

Ethanol exposure: flies were housed in groups of 10 for 3 days as described above. Flies were then exposed to either ethanol (test) or water (control), for 4 consecutive days as described previously by^110^. Flies were then inserted into Fly Bowl arenas for video recording, as described above.

Circadian time shift: flies were housed in groups of 10 for 3 days as described above, or with a two hour time shift (late wake). Flies were then inserted into FlyBowl arenas as housed or as a mixed group of 5 flies from each condition (mixed).

Starvation: flies were collected in groups of 10 as described above. 24 hrs before the behavioral te st, flies were either moved into vials containing agar (starved) or kept in vials with food (controls). Flies were then inserted into Fly Bowl arenas for video recording, as described above.

Ratios of sub populations within a group: WT flies were housed in groups of 10 as described above. Cyp6a20-Gal-4/+; UAS-Cyp6a20-RNAi/+ flies were collected and housed in isolation, as described above for WT isolated flies. Flies were then inserted into FlyBowl arenas in groups of 10, composed of varying amounts of knock-down flies (1 to 5) and WT flies (9 to 5) for video recording. Video recording was performed as described above.

### Statistical analysis

For each experiment except experiments with Cyp6a20 RNAi flies, Shapiro–Wilk test was done on each experiment to test for normal distribution.

For experiments with two-conditions: statistical significance was determined by t-test for experiments that were distributed normally, and by Wilcoxon test for experiments that were not distributed normally.

For experiments with three or four conditions: statistical significance determined by one-way ANOVA followed by Tukey’s range test for experiments that were distributed normally, and by Kruskal–Wallis test followed by Wilcoxon signed-rank test for experiments that were not distributed normally.

Variance: F-test of the equality of two variances was used for all-flies analysis and between-group analysis. Students t-test was used for averages of within groups analysis. FDR correction for multiple testing was performed for all analyses.

Ratios of sub populations normalized to controls: To compare log-ratios of means (test/control), all values were log2-transformed and differences between mean log-values were tested. Specifically, the effect of treatment and mutant number on the fraction of each parameter was tested with a linear regression and a 2-way ANOVA was performed on the resulting model. Log-ratios between different number of mutants were compared in terms of difference of differences defined by linear contrasts and FDR correction was applied to all comparisons.

t-SNE analysis: Visualized using t-Stochastic Neighbor Embedding (t-SNE), using the Barnes-Hut algorithm and implementation (http://homepage.tudelft.nl/19j49/t-SNE.html).

---

**Table S1:**
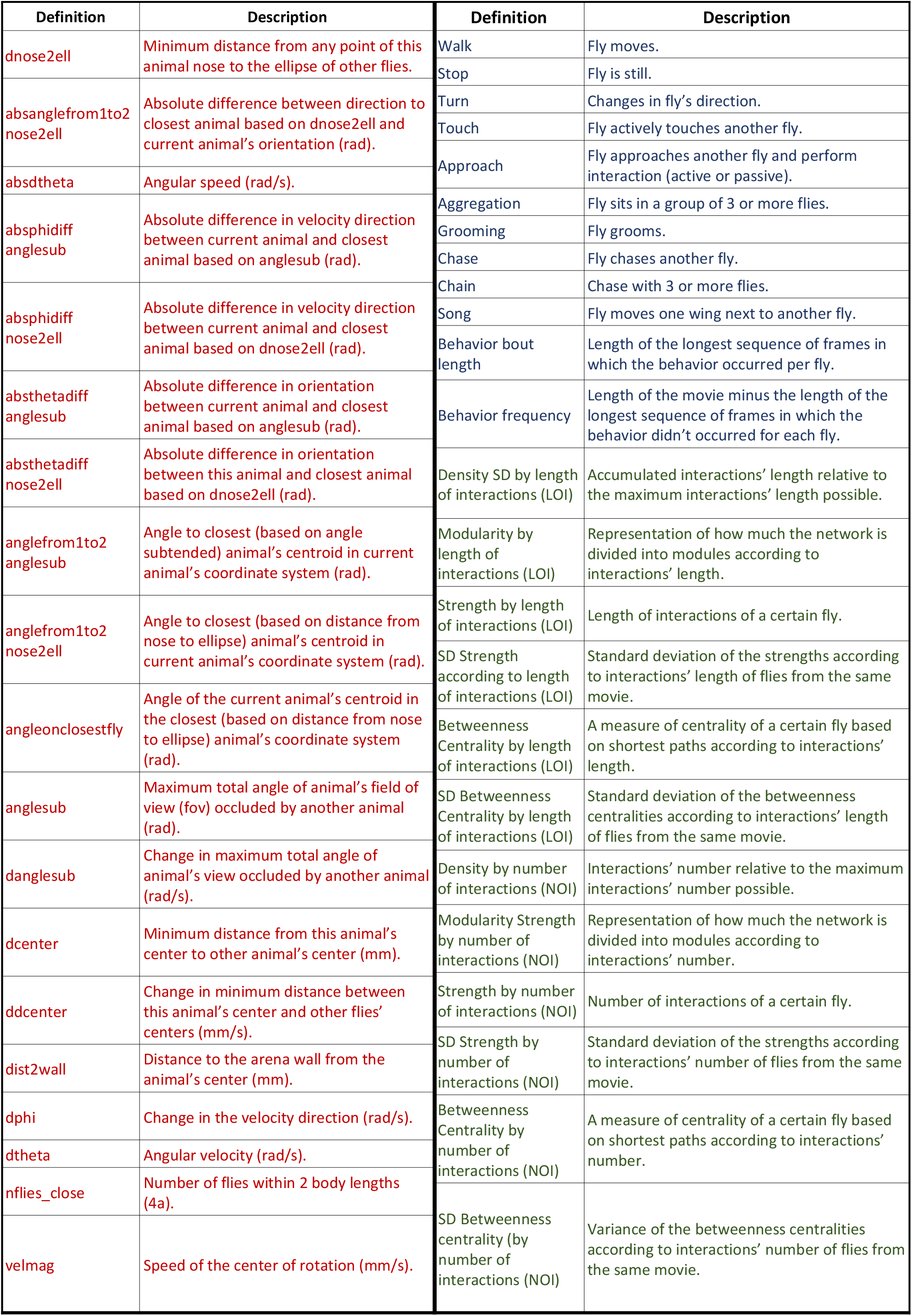
Definitions of behavioral features used in this work. Kinetic (red) features were obtained from Kabra et al. Classified behavioral features (blue) were generated using JAABA. Network (green) features were calculated in R

| Definition | Description | Definition | Description |
| --- | --- | --- | --- |
| dnose2ell | Minimum distance from any point of this animal nose to the ellipse of other flies. | Walk | Fly moves. |
| absanglefrom1to2<br>nose2ell | Absolute difference between direction to closest animal based on dnose2ell and current animal's orientation (rad). | Stop | Fly is still. |
| absdtheta | Angular speed (rad/s). | Turn | Changes in fly's direction. |
| absphidiff<br>anglesub | Absolute difference in velocity direction between current animal and closest animal based on anglesub (rad). | Touch | Fly actively touches another fly. |
| absphidiff<br>nose2ell | Absolute difference in velocity direction between current animal and closest animal based on dnose2ell (rad). | Approach | Fly approaches another fly and perform interaction (active or passive). |
| absthetadiff<br>anglesub | Absolute difference in orientation between current animal and closest animal based on anglesub (rad). | Aggregation | Fly sits in a group of 3 or more flies. |
| absthetadiff<br>nose2ell | Absolute difference in orientation between this animal and closest animal based on dnose2ell (rad). | Grooming | Fly grooms. |
| anglefrom1to2<br>anglesub | Angle to closest (based on angle subtended) animal's centroid in current animal's coordinate system (rad). | Chase | Fly chases another fly. |
| anglefrom1to2<br>nose2ell | Angle to closest (based on distance from nose to ellipse) animal's centroid in current animal's coordinate system (rad). | Chain | Chase with 3 or more flies. |
| angleonclosestfly | Angle of the current animal's centroid in the closest (based on distance from nose to ellipse) animal's coordinate system (rad). | Song | Fly moves one wing next to another fly. |
| anglesub | Maximum total angle of animal's field of view (fov) occluded by another animal (rad). | Behavior bout length | Length of the longest sequence of frames in which the behavior occurred per fly. |
| danglesub | Change in maximum total angle of animal's view occluded by another animal (rad/s). | Behavior frequency | Length of the movie minus the length of the longest sequence of frames in which the behavior didn't occurred for each fly. |
| dcenter | Minimum distance from this animal's center to other animal's center (mm). | Density SD by length of interactions (LOI) | Accumulated interactions' length relative to the maximum interactions' length possible. |
| ddcenter | Change in minimum distance between this animal's center and other flies' centers (mm/s). | Modularity by length of interactions (LOI) | Representation of how much the network is divided into modules according to interactions' length. |
| dist2wall | Distance to the arena wall from the animal's center (mm). | Strength by length of interactions (LOI) | Length of interactions of a certain fly. |
| dphi | Change in the velocity direction (rad/s). | SD Strength according to length of interactions (LOI) | Standard deviation of the strengths according to interactions' length of flies from the same movie. |
| dtheta | Angular velocity (rad/s). | Betweenness Centrality by length of interactions (LOI) | A measure of centrality of a certain fly based on shortest paths according to interactions' length. |
| nflies_close | Number of flies within 2 body lengths (4a). | SD Betweenness Centrality by length of interactions (LOI) | Standard deviation of the betweenness centralities according to interactions' length of flies from the same movie. |
| velmag | Speed of the center of rotation (mm/s). | Density by number of interactions (NOI) | Interactions' number relative to the maximum interactions' number possible. |
|  |  | Modularity Strength by number of interactions (NOI) | Representation of how much the network is divided into modules according to interactions' number. |
|  |  | Strength by number of interactions (NOI) | Number of interactions of a certain fly. |
|  |  | SD Strength by number of interactions (NOI) | Standard deviation of the strengths according to interactions' number of flies from the same movie. |
|  |  | Betweenness Centrality by number of interactions (NOI) | A measure of centrality of a certain fly based on shortest paths according to interactions' number. |
|  |  | SD Betweenness centrality (by number of interactions (NOI) | Variance of the betweenness centralities according to interactions' number of flies from the same movie. |

**Figure S1.**
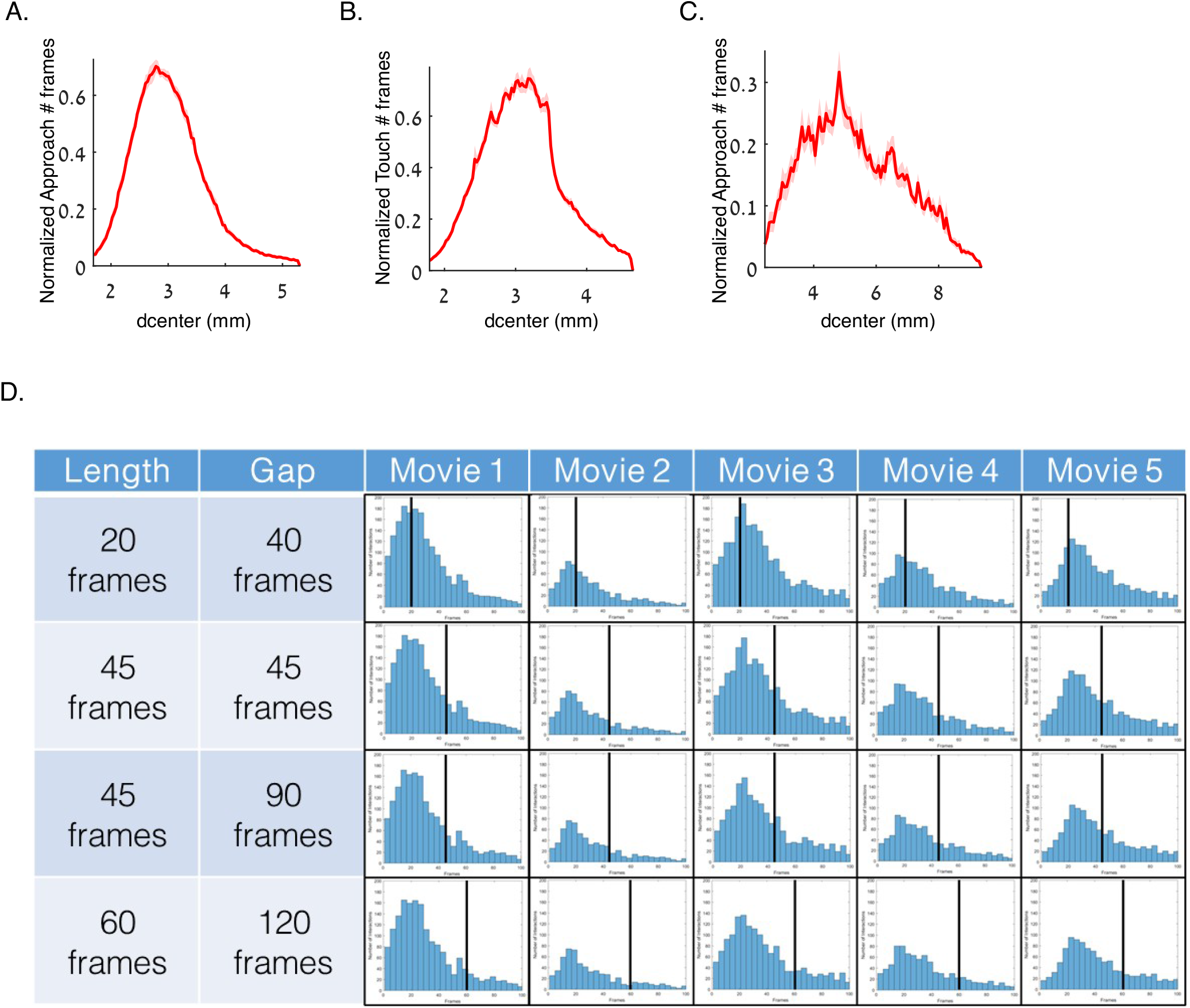
Defining interaction distance, duration and gap thresholds affects the number of very short interactions. A-C. Distribution of normalized number of frames according to the distance between the fly center to another fly center (dcenter) in which approach (A), touch (B) and social clustering (C) as quantified using JAABA. Light red signifies SE. N= 39. D. Number of encounters that meet the minimal distance and angle requirements for interaction as a function of encounter duration in five movies, with different combinations of duration and gap parameters (20-60 frames/0.4-2 sec and 40-120 frames/1.2-4 sec respectively). Each row represents a combination of duration and gap values. Each column represents one movie. Black lines represent the minimal threshold for an interaction according to the specific duration parameter.

**Figure S2.**
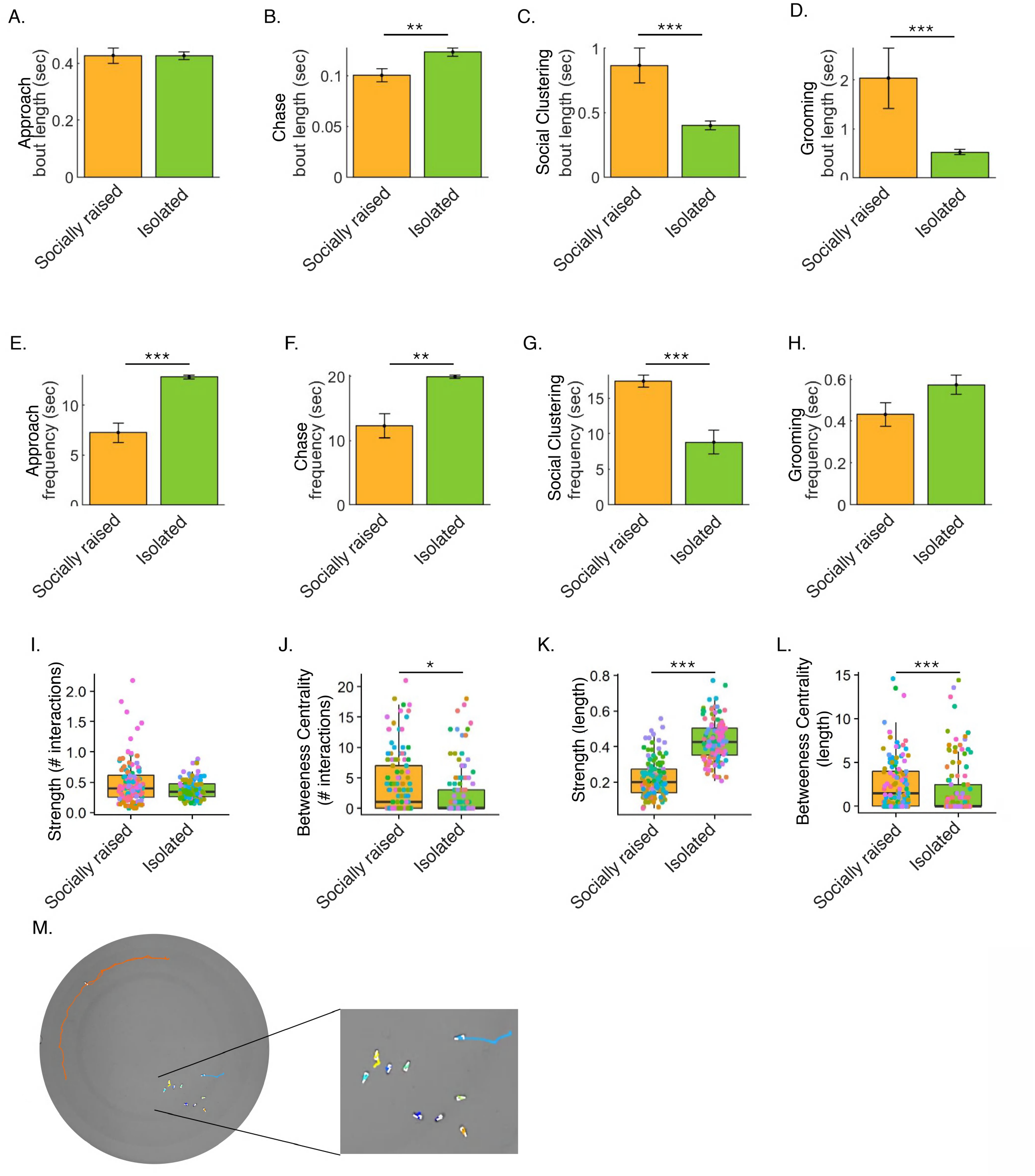
Prior social experience affects bout-length and frequency of specific behaviors and changes network structure. A-H: Average bout-length (A-D) and frequency (E-H) of specific behaviors (Interaction, Chase, Social clustering and Grooming, respectively) of socially raised (orange) vs. isolated (green) WT male flies. I-L: Per-fly network features (Strength and Betweenness Centrality) in which network weights were calculated according to duration of interactions (I-J) or number of interactions (K-L) between socially raised (orange) vs isolated (green) WT male flies. t-test for normally distributed features or Wilcoxon test for non-normally distributed features. FDR correction was applied to all comparisons. N=18, * P<0.05, ** P<0.01, *** P<0.001. Error bars signify SEM. M: Picture of a social clustering event, performed by socially raised WT male flies within a FlyBowl arena, colored lined represent tracking trajectories over the next 60 frames/2 sec.

**Figure S3.**
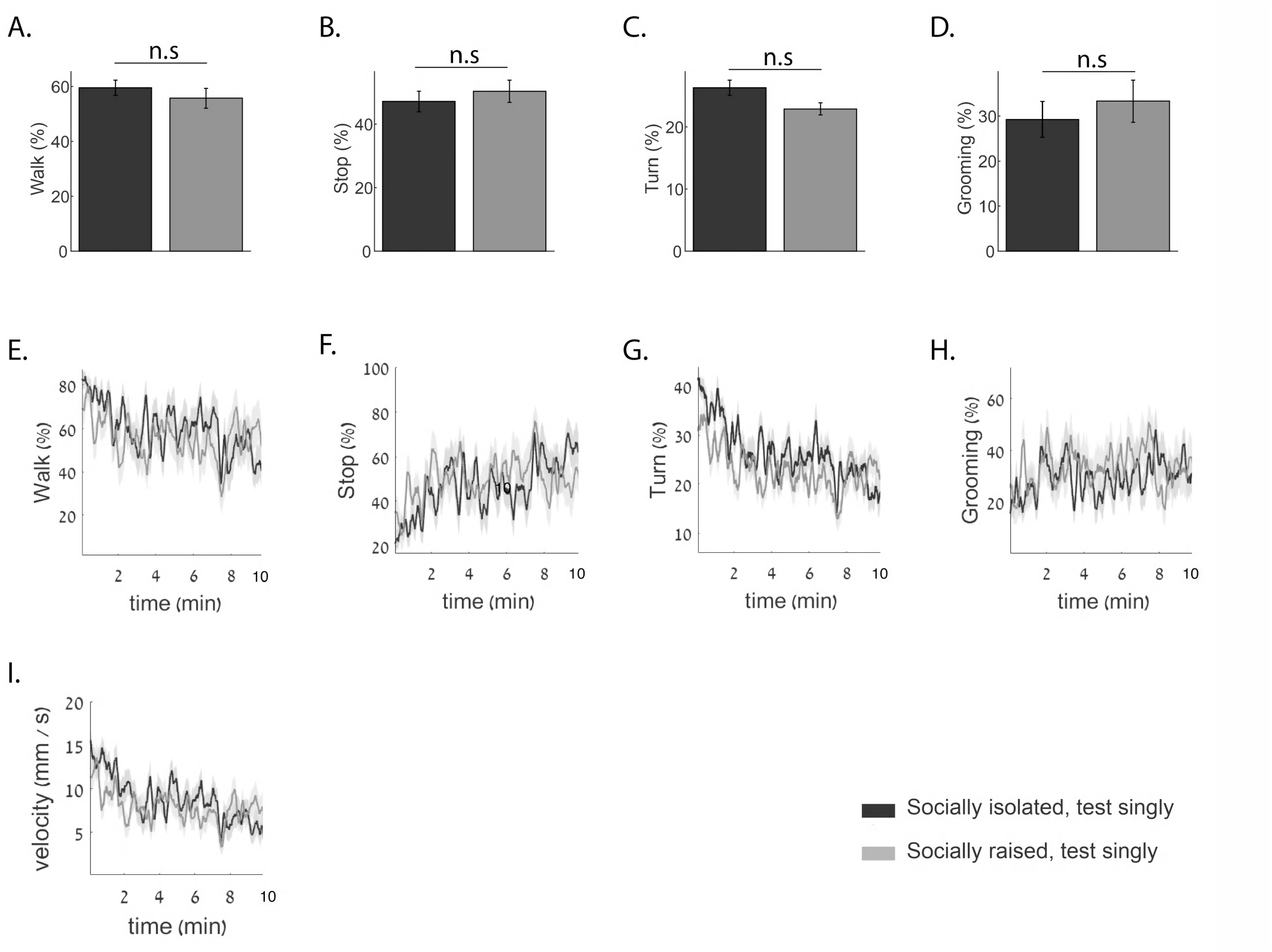
Prior social experience does not affect the behavior of flies tested singly. A-H: Average percentage of time and per-frame averages of previously isolated male flies (black) vs. socially raised male flies (gray) that perform walk (A, E), stop (B, F), turn (C, G) and Grooming (D, H) behaviors. I. average velocity per-frame of previously isolated male flies (black) vs. socially raised male flies (gray). Wilcoxon test P>0.05 N= 17.

**Figure S4.**
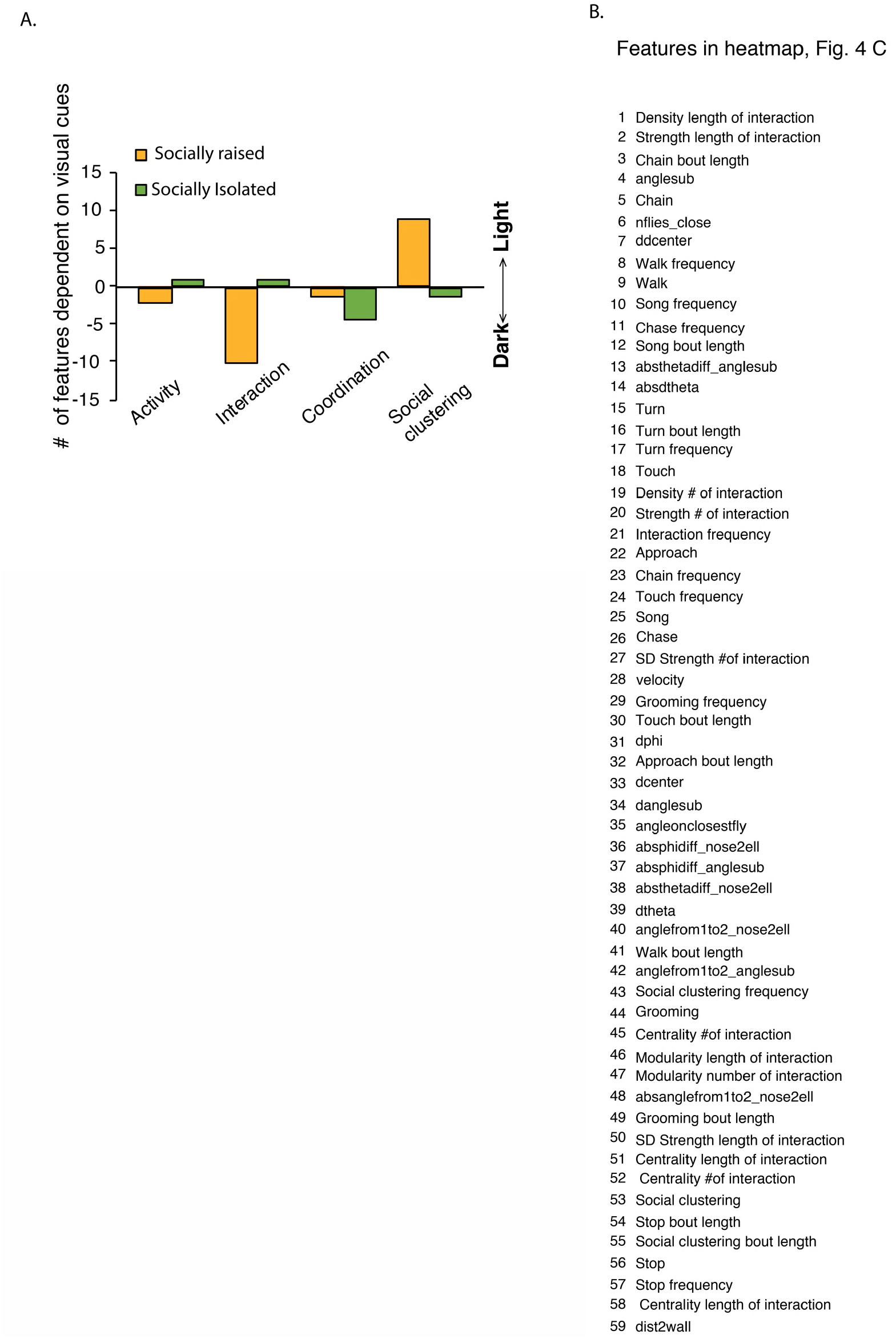
The behavioral signature of socially raised flies displays a higher dependency on visual cues. A. Number of behavioral features that display significantly higher scores in either dark (negative y axis) or light (positive y axis) per condition (isolated-green or raised-orange), divided into 4 categories; activity, interaction, coordination and social clustering related features. B. list of corresponding behavioral features from Fig. 4C.

**Figure S5.**
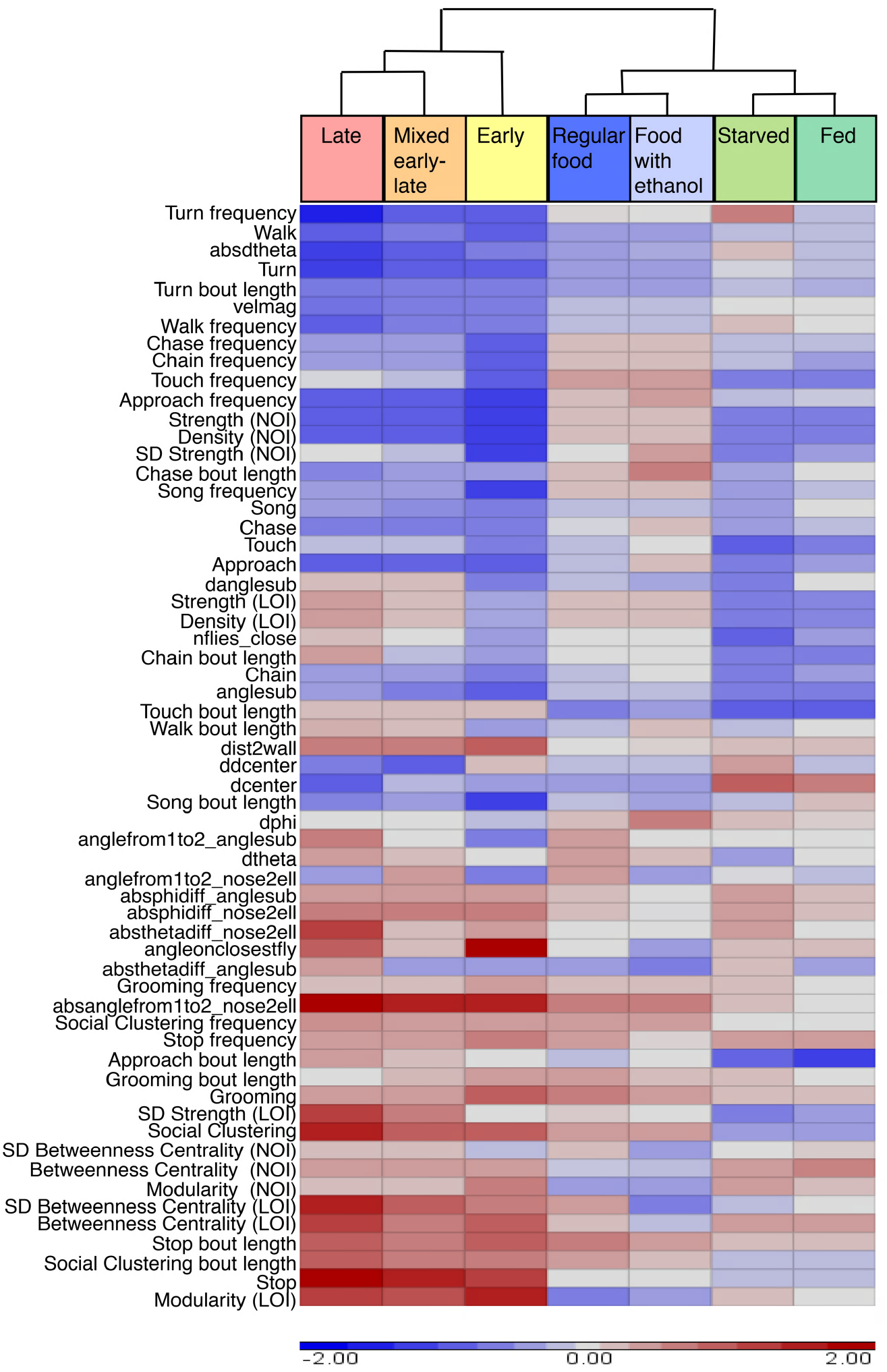
Different internal motivational states in socially raised flies do not affect group signatures. Socially raised flies which were starved for 24 hours prior to behavioral test (fed/starved), exposed to ethanol for 3 days prior to behavioral test (food with ethanol/regular food) or tested at different times during the day (early/mixed/late) do not display any differences in group behavior, compared with controls. Hierarchical clustering of conditions reveals a similarity between each experimental group and its control group (left, hierarchy tree). LOI - calculated according to the length of interactions. NOI - calculated according to the number of interactions. t-test for normally distributed parameter or Wilcoxon test for non-normally distributed parameters in starvation and ethanol experiments. One-way ANOVA with Bonferroni post hoc test for normally distributed parameters or Kruskal Wallis followed by Wilcoxon signed-rank test for non-normally distributed parameters in different times experiment. FDR correction was applied to all comparisons. N=13, 14 and 6 for ethanol, starvation and time difference tests respectively, P>0.05, n.s.

**Figure S6.**
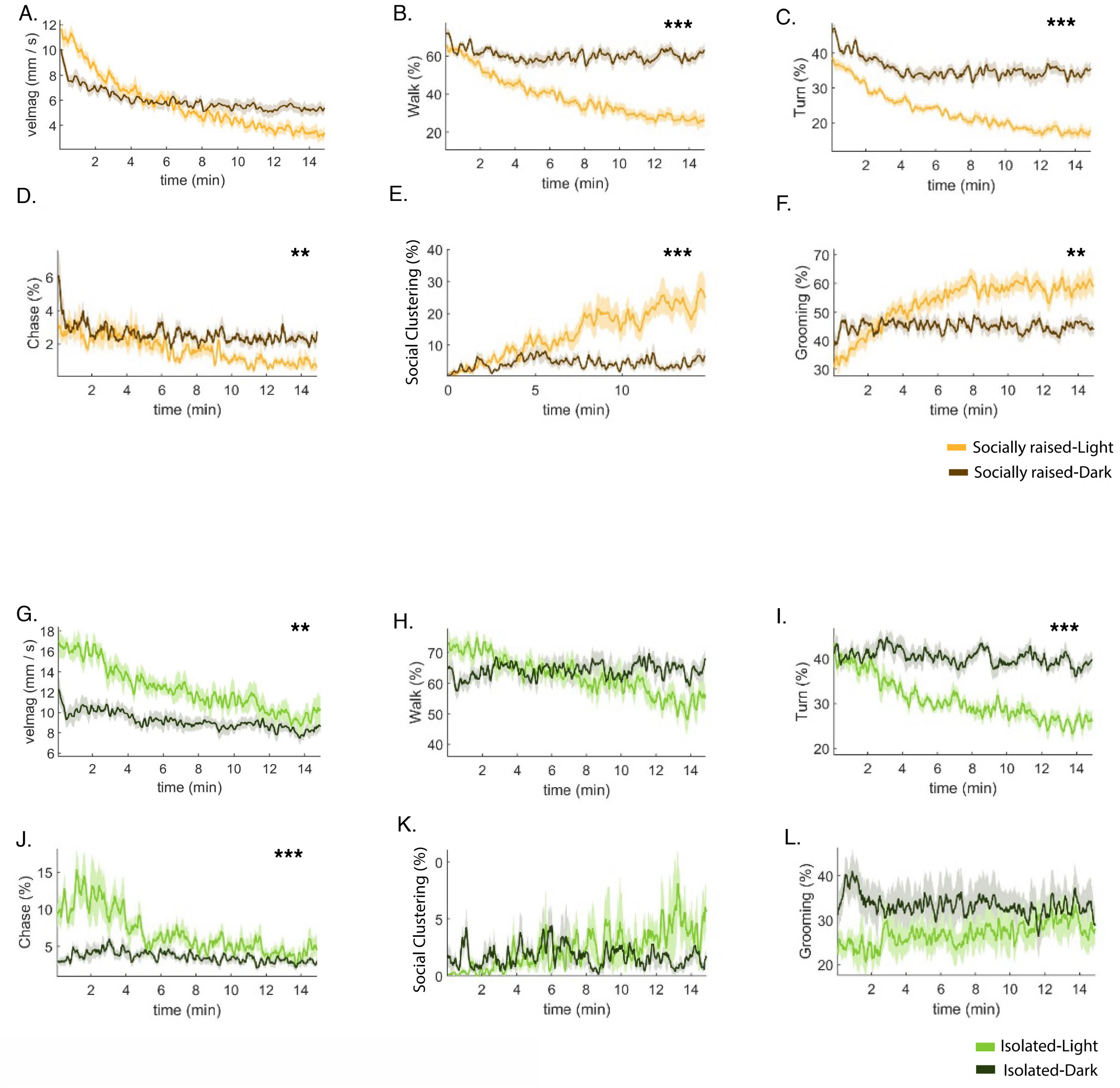
Visual cues are required for habituation during test. A-F: Average per-frame of velocity (velmag), walk, turn, chase, social clustering and grooming of socially raised WT male flies tested in normal lighting (raised light - orange) or in the dark (raised dark - brown). G-L: Average per-frame of velocity (velmag), walk, turn, chase, social clustering and grooming of socially isolated WT male flies tested in normal lighting or in the dark. Statistical analysis was performed on the average of each behavior for the entire duration of the test (15 min). t-test for normally distributed features or Wilcoxon test for non-normally distributed features. FDR correction for multiple testing was performed for all analyses. N=18 for raised and N=11 for isolated experiments, ** P<0.01, *** P<0.001. Error bars signify SEM.

**Figure S7.**
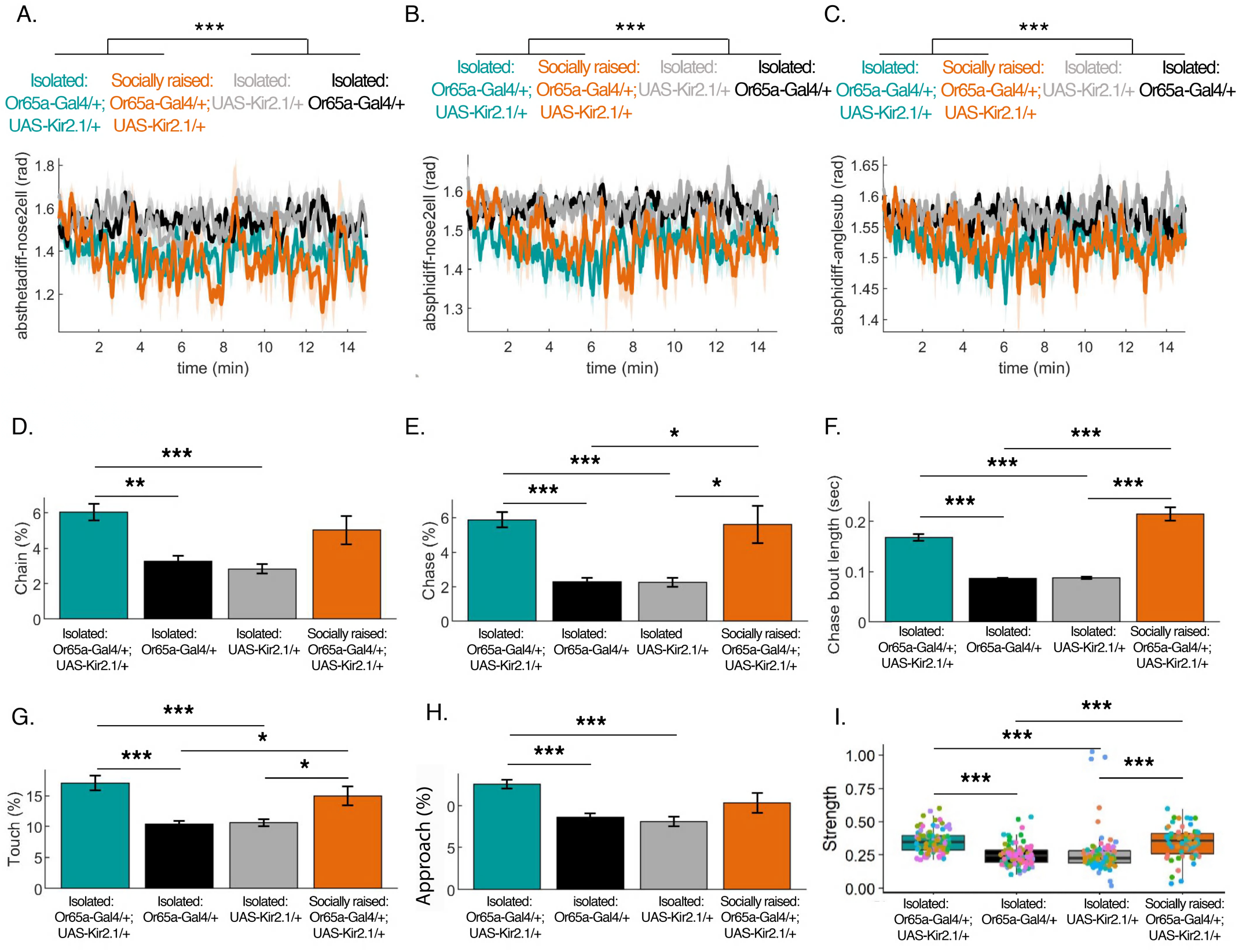
Socially raised and isolated Or65a-Gal4/+; UAS-Kir2.1/+ male flies display similarities in group behavior compared to isolated genetic controls. A-C: Per-frame averages of three kinetic features (A: absthetadiff_nose2ell, B: absphidiff_nose2ell, C: absphidiff_anglesub) in socially raised (orange) and isolated (blue) Or65a-Gal4/+; UAS-Kir2.1/+ flies compared to socially isolated genetic controls (black and gray). D-H: Average percentage of time socially raised (orange) and isolated (blue) Or65a-Gal4/+; UAS-Kir2.1/+ male flies performed chain, chase, chase bout length, touch and interaction behaviors compared with socially isolated genetic controls (black and gray). I: Per-fly network strength of socially raised (orange) and isolated (blue) Or65a-Gal4/+; UAS-Kir2.1/+ male flies compared to socially isolated genetic controls (black and gray). One-way ANOVA with Bonferroni post hoc test for normally distributed parameters or Kruskal Wallis followed by Wilcoxon signed-rank test for non-normally distributed parameters. FDR correction for multiple testing was performed for all analyses. N=6, * P<0.05, ** P<0.01, *** P<0.001. Error bars signify SEM.

**Figure S8.**
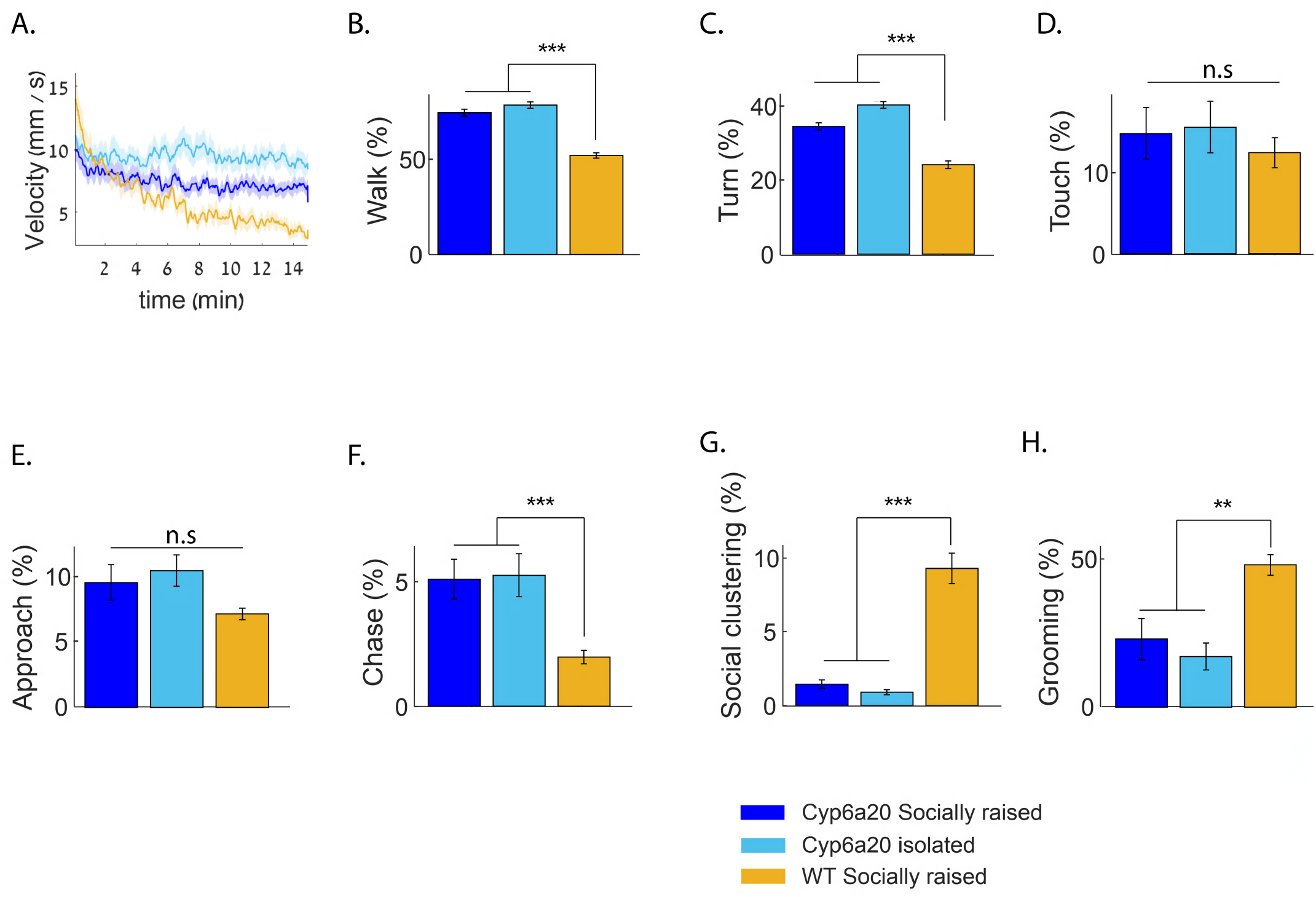
Cyp6a20 knock-down eliminates the effect of social experience on behavior in a group. A. Average velocity per frame of groups composed of 10 socially raised (blue) or isolated (light blue) Cyp6a20-Gal4/+; UAS-Cyp6a20-RNAi flies compared with 10 WT socially raised (orange) flies. B-H. Average percentage of time socially raised vs. isolated Cyp6a20-Gal4/+; UAS-Cyp6a20-RNAi and compared with 10 WT socially raised flies spent in walk (B), turn (C), touch (D), approach (E), chase (F), social clustering (G) and grooming (H) behaviors. Wilcoxon test. N=9. **P<0.01, ***P<0.001.

**Figure S9.**
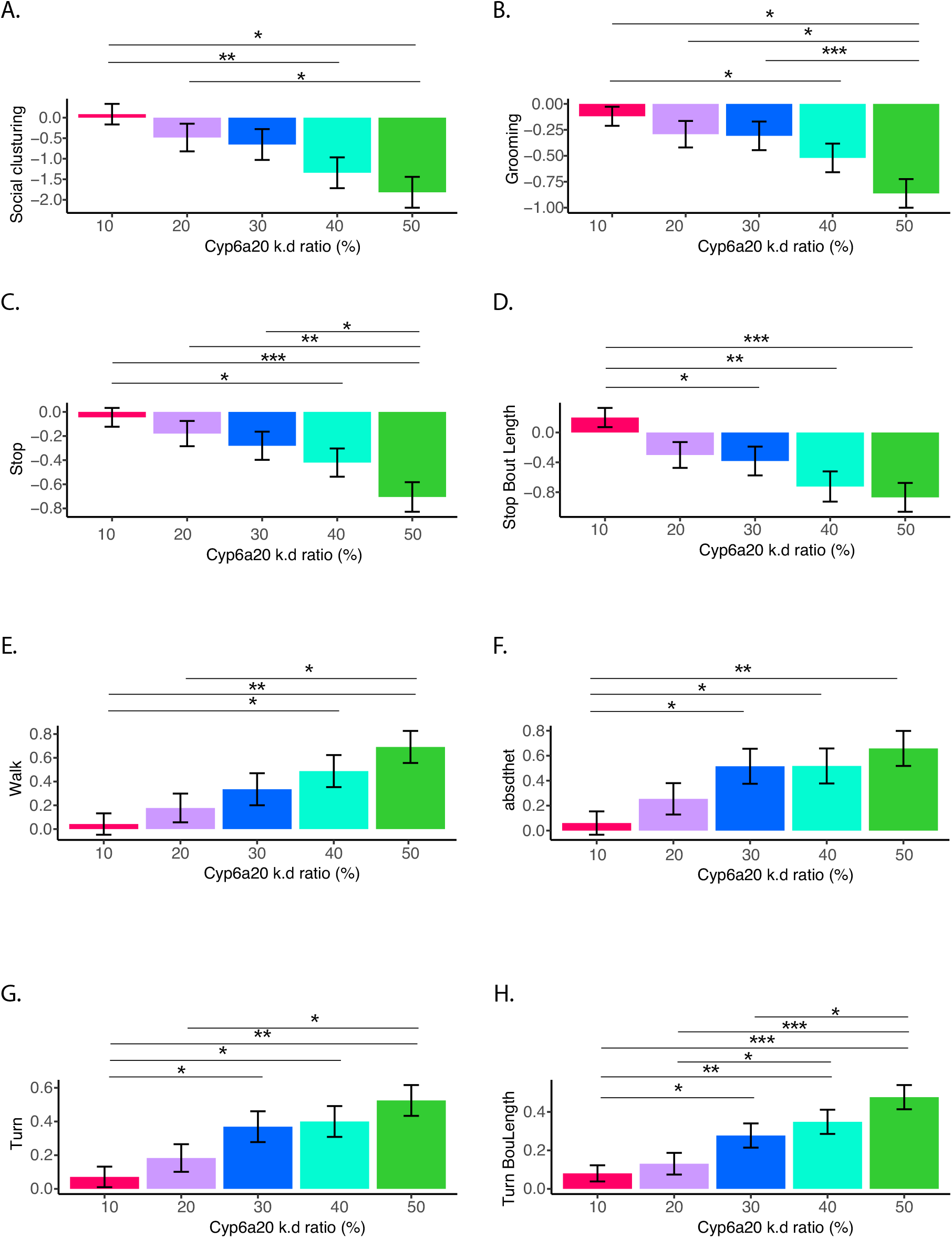
Sub populations in a group affect specific features within behavioral group signatures. A-H: log2 transformed averages of gradually decreasing behavioral features (A: Social clustering, B: Grooming, C: Stop, D: Stop bout length) and gradually increasing features (E: Walk, F: absdtheta, G: Turn, H: Turn bout length) in groups composed of 10%-50% isolated Cyp6a20-Gal-4/+; UAS-Cyp6a20-RNAi to socially raised WT flies. To compare log-ratios of means (test/control), all values were log2-transformed and differences between mean log-values were tested. Specifically, the effect of treatment and mutant number on the fraction of each parameter was tested with a linear regression and a 2-way ANOVA was performed on the resulting model. Log-ratios between different number of mutants were compared in terms of difference of differences defined with by linear contrasts and FDR correction was applied to all comparisons. N=14, 8, and 6 for groups of 10%, 20-30% and 40-50%, respectively, * P<0.05, ** P<0.01, *** P<0.001. Error bars signify SEM.

## Notes

### Competing Interest Statement

The authors have declared no competing interest.

### Summary of Updates

Living in a group creates a complex and dynamic environment in which the behavior of the individual is influenced by and affects the behavior of others. Although social interactions and group living are fundamental adaptations exhibited by many organisms, relatively little is known about how prior social experience, internal states and group composition shape behavior in a group, and the neuronal and molecular mechanisms that mediate it. Here we present a practical framework for studying the interplay between social experience and group interaction in Drosophila melanogaster and show that the structure of social networks and group interactions are sensitive to group composition and individuals social experience. We simplified the complexity of interactions in a group using a series of experiments in which we controlled the social experience and motivational states of individuals to dissect patterns that represent distinct structures and behavioral responses of groups under different social conditions. Using high-resolution data capture, machine learning and graph theory, we analyzed 60 distinct behavioral and social network features, generating a comprehensive representation (group signature) for each condition. We show that social enrichment promotes the formation of a distinct group structure that is characterized by high network modularity, high inter-individual and inter-group variance, high inter-individual coordination, and stable social clusters. Using environmental and genetic manipulations, we show that this structure requires visual and pheromonal cues, and that cVA sensing neurons are necessary for the expression of different aspects of social interaction in a group. Finally, we explored the formation of group behavior and structure in heterogenous groups composed of flies with distinct internal states, and discovered evidence suggesting that group structure and dynamics reflect a level of complexity that cannot be explained as a simple average of the individuals that constitute it. Our results demonstrate that fruit flies exhibit complex and dynamic social structures that are modulated by the experience and composition of different individuals within the group. This paves the path for using simple model organisms to dissect the neurobiology of behavior in complex social environments.

